# LEVERAGING CITIZEN SCIENCE TO ASSESS RICHNESS, DIVERSITY, AND ABUNDANCE IN ANT COMMUNITIES

**DOI:** 10.1101/2024.06.11.598431

**Authors:** Tim M. Szewczyk, Guillaume Lavanchy, Anne Freitag, Aline Dépraz, Amaury Avril, Olivier Broennimann, Antoine Guisan, Cleo Bertelsmeier, Tanja Schwander

## Abstract

Citizen science is a key resource in overcoming the logistical challenges of monitoring biodiversity. While datasets collected by groups of volunteers typically have biases, recent methodological and technological advances provide approaches for accounting for such biases, particularly in the context of modelling species distributions and diversity. Specifically, data integration techniques allow for the combination of scientifically collected datasets with haphazardly sampled presence-only datasets created by most citizen science initiatives. Here, we use a hierarchical Bayesian framework to integrate a set of ant presences collected by citizen scientists in the Vaud canton (Switzerland) with ant colony density data collected concurrently in the same region following a scientific sampling design. The community-level Poisson point process model included species-specific responses to the local (1.2 m^2^) and regional (1 km^2^) environment, with the presence-only samples incorporated at the regional scale to predict local and regional ant communities. At the regional scale, species richness followed a hump-shaped pattern and peaked near 1000 m while abundance increased with elevation. Low elevation and montane ant communities were composed of distinct species assemblages. At the local scale, the link between elevation and richness, diversity, and abundance was weak. At low elevations, local plots varied both in total abundance and species composition, while at higher elevations, the species composition was less variable. The citizen science dataset showed a general tendency toward under-representation of certain species, and heavy spatial sampling bias. Nonetheless, the inclusion of the citizen science data improved predictions of local communities, and also reduced susceptibility to over-fitting. Additionally, the citizen science dataset included many rare species not detected in the structured abundance dataset. The model described here illustrates a framework for capitalizing on the efforts of citizen scientists to better understand the patterns and distribution of biodiversity.

## Introduction

Citizen science is a potential solution to many of the logistical complications that hinder ecological research at broad scales. The many volunteer collectors can sample ecological systems across regional, continental, and even global extents (Theobald et al. 2015), and include private land that is otherwise inaccessible to researchers (Pernat et al. 2020). Accordingly, citizen science initiatives have become increasingly common, with applications ranging from monitoring the spread of invasive rabbits in Australia (Roy-Dufresne et al. 2019) to identifying impacts of hunting on game species’ life history traits in the Amazon (El Bizri et al. 2020). Citizen science could be particularly useful for obtaining information on the abundance and diversity of invertebrate taxa such as insects, since specimens can be collected and easily returned for expert identification or genetic analysis (Ward 2014; Pernat et al. 2020).

Reliable monitoring of insect biomass and diversity is crucial, given that they represent the most numerous group of animals on the planet with an estimated 5.5 billion species (Stork 2018), play key ecological roles and provide many ecosystem services. Over the past decades, global declines of insects have raised concerns, with for example an estimated decline of 75% of flying insects over 30 years (Hallmann et al. 2017). Given the enormous task of assessing richness, diversity and abundance in insect communities, the efforts of citizen scientists represent a valuable resource.

A potential problem with citizen-science datasets however is that the decentralized and unstructured nature of the data collection typically results in sampling bias due to differences in species detectability and sampling effort in space (e.g., Theobald et al. 2015; Steen, Elphick, and Tingley 2019; Duan et al. 2020; Henckel et al. 2020). To use the full potential of citizen-science datasets, it is therefore crucial to account for such bias (e.g., Isaac et al. 2014; Steen, Elphick, and Tingley 2019; Johnston et al. 2020; Robinson et al. 2020). A powerful way to correct for various sources of bias in citizen-science datasets is to combine these datasets with structured samples collected by researchers using data-integration techniques (Isaac et al. 2019; Miller et al. 2019). In addition to maximizing the number of observations used, data-integration techniques aim to exploit the strengths of each dataset. For example, presence-only data often detect more rare species, while datasets collected in a structured manner better represent community composition and abundance (Steen, Elphick, and Tingley 2019; Henckel et al. 2020; Pernat et al. 2020). Combining large-scale citizen science approaches with small-scale structured species inventories could thus be an ideal solution for long-term monitoring of insect biodiversity and abundance.

Here, we provide a unique dataset on ants from the Vaud canton (Switzerland) to assess the usability of citizen-science datasets for insect monitoring and the value of complementing citizen-science datasets with structured samples. Our dataset combines presences of ants collected in a citizen science project (Avril et al. 2019; Freitag et al. 2020) with local abundance data from a concurrent structured sampling effort. We develop a community-level Poisson Point Process Model (PPM) within a hierarchical Bayesian framework, to jointly model local ant colony density (structured sampling) with additional presence-only information (citizen science data) integrated at a regional scale, while accounting for taxonomy and correlated residuals among species resulting from unmeasured variables such as sampling bias or biotic interactions. We use this model to predict the patterns of ant colony density, richness, diversity, and community structure across the landscape at local and regional scales while incorporating uncertainty in species compositions, and to evaluate the strength of selected environmental drivers across spatial scales. We compare inferences from the integrated model with those from models built on the structured dataset only, and further assess the differences in observed communities from each sampling method. We show how integrating scientific and citizen-science inventory data allows capitalizing on the efforts of citizen scientists to better understand the patterns and distribution of biodiversity.

## Methods

### Study region & sampling design

This study focuses on ant species in the Vaud canton in western Switzerland (46.2–47.0°N, 6.1– 7.2°E), a topographically heterogeneous region (372–3210 m) composed of the Swiss plateau in the center, the Jura mountains in the west, and the Alps in the east (Fig. 1). The hilly central plateau is dominated by anthropogenic land use, including cropland, vineyards, pastures, and urban zones, with patches of predominantly deciduous forest. The mountain ranges rise steeply from approximately 1000m and span the forested montane and subalpine zones through the alpine zone, with widespread cattle grazing (Delarze et al. 2015; Gago-Silva, Ray, and Lehmann 2017; Beck, Rüdlinger, and McCain 2017).

**Figure 1.**
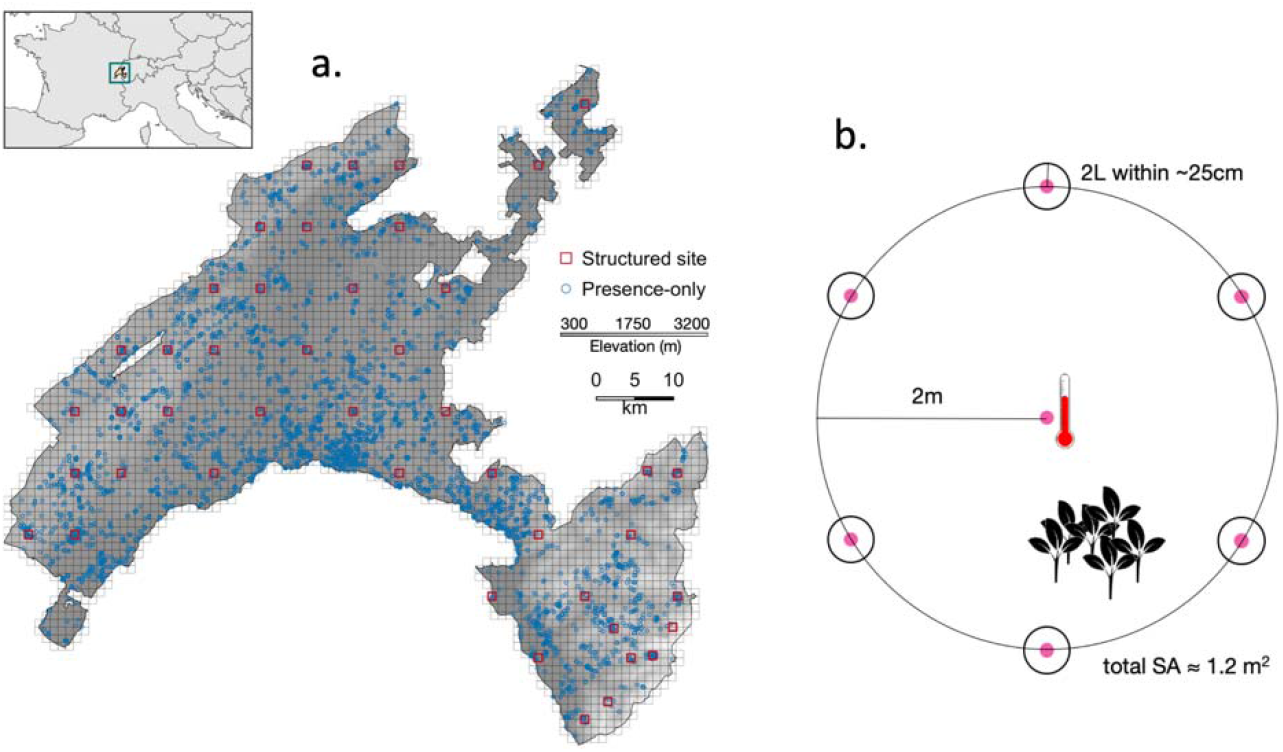
Map of the sampling design in Vaud, Switzerland. (a) Samples consisted of presence-only data (blue points) aggregated to a 1 km grid (light gray) and structured abundance data collected at 1 km long-term biodiversity monitoring sites (red squares). (b) Sampling plot design within structured sites.

Ants were collected during the summer of 2019 in two simultaneous efforts (Fig. 1), comprising a citizen science presence-only dataset and a structured abundance dataset. The citizen science project (“Opération Fourmis”; Avril et al. 2019; Freitag et al. 2020) was organized to survey the ant fauna within the Vaud canton. Briefly, vials of ethanol were distributed to interested participants, who were encouraged to explore under rocks, on bark, inside twigs, and in downed wood. The protocol asked to collect approximately 10 ant workers from a single colony per vial. An online map was updated periodically to informally highlight data-sparse areas. Collectors returned the vials along with the collection date and the locality of the sample, including geolocation and a brief habitat description. For these presence-only data, we discretized the landscape into a 1 km^2^ grid (3,558 km^2^) and tallied the number of occurrences for each species in each cell.

For the structured abundance dataset, local ant densities were estimated within 44 sites (1 km^2^ each), with standardized effort across sites. The sites were arranged on a regular grid with approximately 5-7 km between adjacent sites. At each site, ant densities were characterized by 25 plots, distributed among 15 habitat types in approximate proportion to the abundance of each habitat, where each habitat present within the site was represented by at least one plot (Supplemental Table 1; see also Freitag et al. 2020). Inaccessible areas (e.g., cliffs, water, property beyond Vaud) were excluded, resulting in several sites with areas less than 1 km^2^, and the number of plots reduced proportionally (1059 total plots).

Each sampling plot consisted of a 2 m radius circle, with soil temperature recorded in the center approximately 6 cm deep, and vegetation characterized according to Braun-Blanquet coverage classes for grass, forb, shrub, litter, bare ground, and moss within the plot (Douglas and Shanholtzer 1978). Six flags were evenly spaced around the circumference. Within ∼25 cm of each flag (total surface area ∼1.2 m^2^), we searched for ant colonies within any downed wood or stumps, under large rocks, and in 2 L of soil, litter, and small rocks using 18 cm Hori Hori gardening knives. We haphazardly collected 10 workers from each colony, placing them directly into vials of ethanol. Within each plot, all trees ≥3 cm diameter at breast height were also inspected for ant workers which were collected regardless of whether or not a nest was identifiable. Lastly, transect lines were mapped *a priori*, distributed proportionally among habitat types and totalling 2 km. Transects were surveyed by walking at approximately 4 km/h, with workers collected from all permanent above-ground mounds within 2 m of the transect line. Because of the differing sampling methods, the tree and transect collections were incorporated into the presence-only dataset.

Thus, the two datasets characterize the regional ant fauna in distinct ways. The presence-only dataset was spatially expansive and included free investigation of subjectively suitable habitats, and was therefore expected to include more rare species. However, collections were also expected to be spatially and taxonomically biased, with more effort near human-dominant areas and toward larger, more obvious, or more anthropophilic species (Ward 2014; Troudet et al. 2017). In contrast, the structured abundance dataset was collected with uniform sampling effort, with a design aimed to produce representative samples of colony density within each site. This dataset was thus expected to better characterize the ground-nesting ant community structure.

All ants were identified to species or species groups based on morphology by taxonomists with all specimens stored at the State Museum of Natural Sciences of Lausanne, Switzerland. Four genera (*Camponotus, Tapinoma, Temnothorax, Tetramorium*) were further identified based on the mitochondrial gene COI due to uncertainties in morphological identifications in these groups. Three legs were taken from one random worker in each sample. DNA extraction, PCR and sequencing for *Camponotus, Tapinoma* and *Temnothorax* was carried out by by the Canadian Center for DNA Barcoding (Guelph, Canada). A fragment of the COI mitochondrial gene was amplified using primers LepF1 and LepR1 (Hebert et al. 2004). The specific protocols used for extraction, PCR amplification and sequencing are available at https://ccdb.ca/resources (last checked 14.07.2022). DNA extraction and COI barcoding of *Tetramorium* was performed by AllGenetics & Biology SL (A Coruña, Spain). The same primers were used for sequencing.

COI sequences of reliably identified individuals were retrieved from published studies and from the Barcode Of Life Database (https://www.boldsystems.org/index.php/Public_BINSearch) and added as references for *Tapinoma* (Seifert et al. 2017), *Temnothorax* (Blatrix et al. 2020) and *Tetramorium* (Wagner et al. 2017). The sequences were aligned with the L-INS-I algorithm of *mafft* 7.481 (Katoh et al. 2017). The phylogeny was constructed with *iq-tree* 2.2.0.5 (Minh et al. 2020). The resulting tree was displayed and analyzed with the package *phytools* 0.7-80 (Revell and Reynolds 2012) in R 4.1.1 (R Core Team 2021). We assigned species names to each mtDNA clade based on the morphological identification of the majority of individuals belonging to that clade. For clades where a significant proportion of individuals were identified morphologically as a differed species than the majority, we used the species names of the reference individuals that grouped with these clades.

### Environmental variables

Environmental variables were selected based on theoretical or empirical support in the literature as likely drivers of ant distributions, richness, and diversity, (Bishop 2017; Liu et al. 2018; Szewczyk and McCain 2019; Longino, Branstetter, and Ward 2019; Uhey et al. 2020). At a regional (1 km^2^) scale, we included growing degree days from ENVIREM as well as its square (Title and Bemmels 2018), annual precipitation from CHELSA (Karger et al. 2017), net primary productivity as the average values across years 2010–2019 (MODIS: MOD17A3), the Shannon diversity of the 15 land cover types based on proportional composition (Gago-Silva, Ray, and Lehmann 2017), the coverage of edge, forest, and agricultural habitats (Gago-Silva, Ray, and Lehmann 2017), the average north-facing aspect calculated as the cosine of the aspect in degrees based on the ASTER digital elevation model (Tachikawa et al. 2011), the log-transformed total length of roads (OpenStreetMap 2019), and the log-transformed total perimeter of buildings (OpenStreetMap 2019). All variables were summarised as the mean or total value within each 1 km^2^ grid cell and within each 1 km^2^ structured site.

Local variables were collected in the field at each structured sampling plot as described above. We included relative soil temperature as a measurement of local variation in temperature, where the recorded temperatures among plots were z-transformed within each site. Canopy was classified as open, closed, or mixed according to land cover type (Supplemental Table 1). Open habitats were categorized as pasture, crop, or other according to field observations. Finally, local productivity was quantified by using the midpoint of the range of percent cover values in each Braun-Blanquet category and summing across grass, forb, and shrub in each plot (Douglas and Shanholtzer 1978; McCain et al. 2018; Szewczyk and McCain 2019).

Variable processing and summarising was performed in R 3.6.3 (R Core Team 2020), with spatial computations in the R packages *raster 3*.*3*.*7* and *sf 0*.*9*.*4*. All variables were z-transformed after summarising to the appropriate spatial scale to improve model behavior (Carpenter et al. 2017).

### Model overview

To integrate the two datasets, we chose Bayesian methods as they naturally and flexibly accommodate the incorporation of multiple datasets (Clark 2005; Beck et al. 2012; Szewczyk and McCain 2019). To model species distributions we used inhomogenous Poisson Point Process Models (PPMs), which model continuous, spatially varying intensities that produce Poisson-distributed occurrences (Renner and Warton 2013; Renner et al. 2015). PPMs are an especially appealing option for combining different datasets. Observations collected with different methods can be modelled using appropriate sampling submodels based on the same underlying intensities (Fithian et al. 2015; Hefley and Hooten 2016; Koshkina et al. 2017; Fletcher et al. 2019; Renner, Louvrier, and Gimenez 2019), and the intensities can be integrated to the appropriate resolution, allowing patterns and processes to be linked across spatial scales (Keil and Jetz 2014; Hefley and Hooten 2016).

Inhomogenous Poisson PPMs assume that the distribution of occurrences is dependent on the variation in local intensity, which may be observed imperfectly resulting in a thinned point process (Warton and Shepherd 2010; Baddeley, Rubak, and Turner 2015; Fithian et al. 2015). One key benefit of PPMs is that the underlying latent intensity is continuous in space, and can be integrated to arbitrary spatial resolutions (Renner et al. 2015; Hefley and Hooten 2016; Koshkina et al. 2017; Fletcher et al. 2019). Following the structure of the sampling design, we modelled the expected intensity of each species at two resolutions (regional: 1 km^2^, local: 1.2 m^2^), representing the area of the sampling sites and the area of the sampling plots respectively. We modelled local intensities as a function of species’ responses to the local and regional environment, with the structured abundance data incorporated at the local scale, and the presence-only data incorporated at the regional scale. The sampling submodel for each data type reflects the collection methods (Isaac et al. 2014; Hefley and Hooten 2016; Fletcher et al. 2019; Miller et al. 2019).

### Model structure

The hierarchical PPM integrates the two datasets, **W** (citizen science data, presence-only; resolution: 1 km) and **Y** (scientific sampling, structured local abundance; 1.2 m plots within 1 km sites) to predict ant communities across the landscape (Fig. 2). Species’ responses to the regional environment are informed by both **W** and **Y**, while responses to the local environment are informed by **Y**. Similarly, **W** helps to identify overall relative abundance among species, while **Y** is a direct measurement of local abundance. We assume that local colony density for each species is a function of local and regional environmental conditions, with potential phylogenetic conservatism among species-specific responses and residual correlation among species.

**Figure 2.**
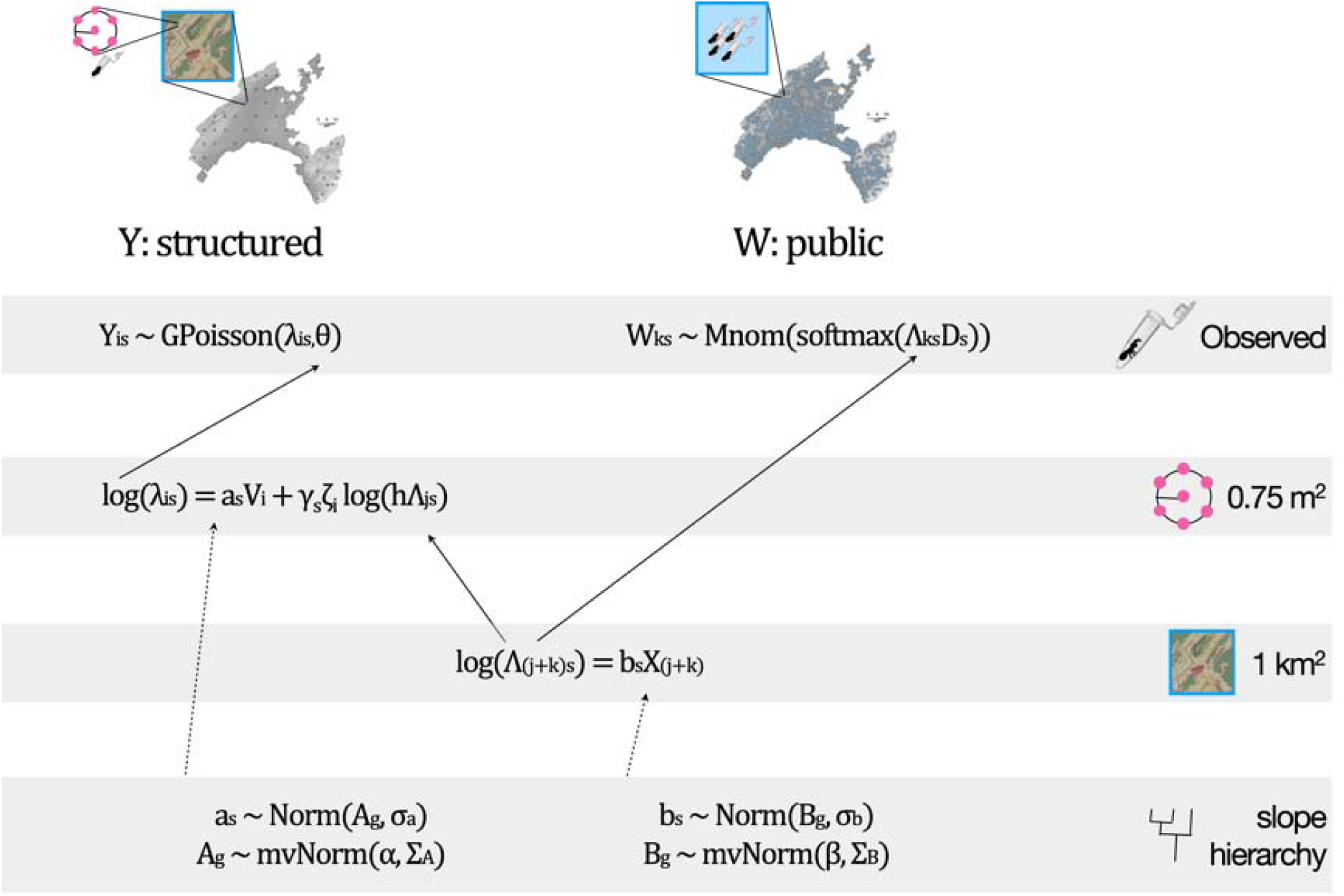
Overview of the model structure. See text for explanations of variables.

The structured abundances, **Y**, are observed at the local scale. For each species *s* = 1 … *S* at plot *i*, the number of observed colonies *Y*_*is*_ follows a generalized Poisson distribution with the latent colony intensity *λ*_*is*_ and dispersion term *θ* to account for overdispersion (Consul and Famoye the 1992; Ntzoufras, Katsis, and Karlis 2005; Isaac et al. 2019; Miller et al. 2019). The local Intensity *λ*_*is*_ is a function of the local environment and the regional intensity of species *s* at encompassing 1 km^2^ site *j*:

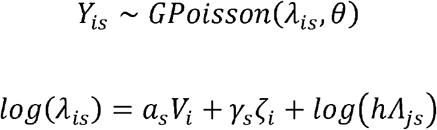

where **V** is a matrix of local environmental covariates, *a*_*s*_ is a vector of species-specific responses, *ζ*_*i*_ is a latent factor representing unmeasured local environmental variables or biotic interactions with *γ*_*s*_ as a species-specific latent factor coefficient constrained between − 1 and 1 (Ovaskainen et al. 2016; Caradima, Schuwirth, and Reichert 2019; Tobler et al. 2019), *h* is a constant scaling factor representing the proportion of site *j* sampled by plot (1.2 m^2^ / 1km^2^ = 1.2e-6), and *Λ*_*js*_ is the regional intensity of species *s* at site *j*. Thus, *log* (*h Λ*_*js*_) functions as a site-level intercept, determining the baseline expected intensity at each plot within a site (Yamaura et al. 2016; Miller et al. 2019).

The presence-only data, **W**, are observed as detections at a regional scale. The detections ofSpecies *s* = 1 … *S* within each 1 km^2^ cell *k* follow a multinomial distribution such that:

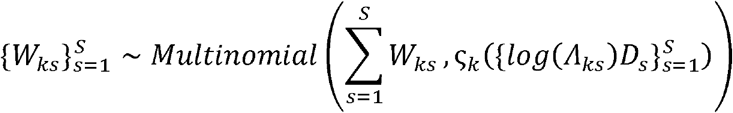

Where 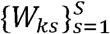 is a vector of length *S* with the number of detections of each species within cell 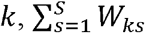 is the total number of detections within the cell, *D*_*s*_ is the proportional detection bias for each species which accounts for taxonomic bias in effort, and 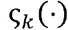is the softmax functionwhich converts the weighted intensities to probabilities that sum to 1 within each cell. Thus, for Species *s* in cell *k*, the observed count *W*_*ks*_ is expected to be higher if the species has high relative abundance (*Λ*_*Ks*_ is larger relative to other species), more samples were collected (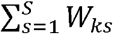 is larger), or if species *s* is over-represented in **W** relative to **Y** (*D*_*s*_ is larger). The presence-only counts thus inform relative patterns across the landscape, while accounting for spatial and taxonomic bias in sampling intensity (Isaac et al. 2014).

While **W** and **Y** are modelled with distinct sampling submodels, both *Λ*_*js*_ and *Λ*_*Ks*_ represent the latent intensity of species *s* at an identical 1 km^2^ regional scale. Consequently, the regional intensity *Λ*_(*j,k*)*s*_ is a shared parameter that links the two datasets (Hefley and Hooten 2016; Isaac et al. 2019; Miller et al. 2019). The ecological processes driving variation in regional intensity are thus independent of sampling method, and the intensities are modelled together in a single regression such that:

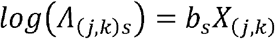

Where *b*_*s*_ is a vector of species-specific slopes, and **X** is a matrix of regional environmental covariates. Thus, the magnitude and spatial variation of the 1 km^2^ intensity *Λ*_(*j,k*)*s*_ is informed by *Y*_*is*_ (Eq. 1,2) with the spatial variation further informed by *W*_*ks*_ (Eq. 3).

The slopes *a*_*s*_ and *b*_*s*_ are species-specific responses at local and regional scales, respectively, and distributed about genus-level means *A*_*g*_ and *B*_*g*_ with standard deviations σ_*a*_ and σ_*b*_. The genus-level means are in turn distributed about aggregate means, *α* and *β*, with correlation are The matrices *∑*_*A*_ and *∑*_*B*_, which reflect the overall responses of the ant community to environmental variables at each resolution while accounting for relatedness at the genus level (Hadfield and Nakagawa 2010; Ovaskainen and Soininen 2011; Szewczyk and McCain 2019; Caradima, Schuwirth, and Reichert 2019).

Several quantities were calculated in each sample from the posterior. Probability of presence was calculated as 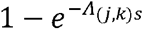, at the regional scale (Hefley and Hooten 2016), and 1 − *pr*(0 ∣ *λ*_*is*_,*θ)* at the local scale. Predicted richness was calculated as the sum of species-level Bernoulli draws based on the probability of presence. Shannon’s H was calculated using *λ*_*i,1:S*_ and *Λ*_(*j,k*)*1:S*_. Total predicted intensity was calculated as 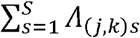 and 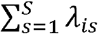. Posterior predictions of local communities were for only structured abundance plots, while regional predictions were for the 44 structured sites and the entire gridded landscape for Vaud (Fig. 1).

### Variable selection & model fitting

To evaluate the effect of including the presence-only data, **W**, we compared a version of the model with and without **W**. We refer to the model that uses both **W** and **Y** as the *Joint* model, and the model using only **Y** as the *Structured* model. The *Structured* model therefore did not include Eq. 3, but nevertheless included species only detected in **W**. We performed variable selection separately on the two models.

Because comparing all possible models was not logistically feasible, variable selection was performed using a 4-fold cross-validation with a step-wise forward search. The 44 sites in **Y** were divided into 4 subsets using the R package *caret 6*.*0*.*86*. The models were parameterized using 3 of the subsets and predictive ability assessed using the 4^th^, with all subsets predicted in turn. For each successive level of model complexity, the expected log pointwise predictive density was calculated for I_*oos*_ out-of-sample plots as 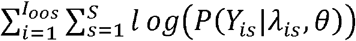 and compared using the R package *loo 2*.*3*.*1*. Variables were added sequentially, and the variable sets for the *Joint* and *Structured* models with the best out-of-sample predictive ability were considered optimal. Using the optimal variable sets, the models were fit using all observations, with intensities then predicted across the full landscape, excluding cells with covariates more than three standard deviations beyond the values available for fitting.

We assessed model performance of the optimal models based on deviance using **Y** such that 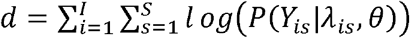 in each sample from the posterior, with the explanatory power of each proposed model as 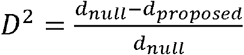(Caradima, Schuwirth, and Reichert 2019; Guisan and Zimmermann 2000). We used an intercept-only model fit with **Y** (i.e., *Structured* model) as the null model. We calculated overall *D*^2^ using all abundances, species-specific 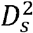 to compare performance across species, and plot-specific 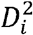 to compare performance across elevations. We calculated each metric for the fitted full models, and during the 4-fold cross validation to assess improvement in predictions of out-of-sample plots.

All models were written in Stan (Carpenter et al. 2017) and run in *CmdStan 2*.*23*.*0*. Prior distributions (Supplemental Table 2) were lightly informative to constrain the sampling algorithm to plausible ranges (Carpenter et al. 2017; Lemoine 2019). During variable selection, we ran 3 chains for each model with 2,500 iterations per chain, with 2,000 iterations as burn-in.For the optimal models, we ran 12 chains with 2,250 iterations each, with 2,000 iterations as warm-up. Model behavior and convergence was assessed by confirming that all 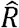 values were ≤1.1, visually inspecting a selection of trace plots including hyperparameters and scale parameters, and confirming that no divergent transitions occurred during the sampling. Highest Posterior Density Intervals (HPDIs) were calculated for all parameters of interest using the R package *HDInterval 0*.*2*.*2*. Code for the full model is available in the supplementary section Code A.1.

### Community analyses

We performed several analyses of the predicted communities with the *λ* posterior medians. We assessed similarity in the taxonomically-weighted community structure among plots with a double principal coordinates analysis (DPCoA) using the R packages *ade4 1*.*7*.*15* and *vegan 2*.*5*.*6* (Dray, Pavoine, and Aguirre de Cárcer 2015; Pavoine 2019). Using relative abundances, the DPCoA accounts for relatedness among species, and we used 100m elevational bins and regions to categorize communities *post-hoc*. The elevational distribution of each species was also characterized by the observed elevational range, and by the proportion of total intensity in low (<1000m) or high elevation (≥ 1000m) environments.

With the *λ* posterior medians, we calculated the *β*-diversity among plots within each site using the R package *betapart 1*.*5*.*1*. The overall *β*-diversity at each site was partitioned into the balanced variation and abundance-gradient components, representing changes in relative abundance among species and changes in total abundance, respectively (Baselga 2017). Change across elevation for total *β*-diversity and each component was assessed using linear and quadratic regressions compared with AICc using the R package *AICcmodavg 2*.*3*.*0*. When *Δ*AICc was ≤4, the linear model was selected as most parsimonious.

### Results

A total of 79 species were detected across both datasets, with 76 in the citizen science presence-only dataset, **W**, and 51 in the structured abundance dataset, **Y** (Supplemental Table 3). **W** contained 6,632 samples collected by citizen scientists and 363 supplemental transect or tree samples, covering 42% of the 1 km^2^ grid (1,309 cells; occurrences per cell: range=1–162, median=3). **Y** contained 1,090 colonies detected across the 44 sites (per-site range=6–58, median=22.5).

The *Joint* model, fit with **W** and **Y**, consistently outperformed the *Structured* model, fit with only **Y**, in predicting novel local communities during cross-validation, and was more resistant to over-fitting (Fig. 3, Supplemental Table 4; optimal models: *Joint D*^2^=0.079; *Structured D*^2^=0.050). In the full model, the *Structured* model fit the data somewhat more closely (*Joint D*^2^ =0.19; *Structured D*^2^=0.21). This same pattern held on average across species (median: *Joint* 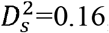*Structured* 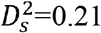) and plots (median: *Joint* 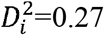; *Structured* 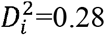). However, the *Joint* model dramatically improved fit for species that were not detected in **Y** (median: *Joint* 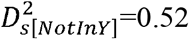 *Structured* 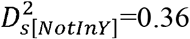). There was no elevational pattern in 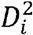.

**Figure 3.**
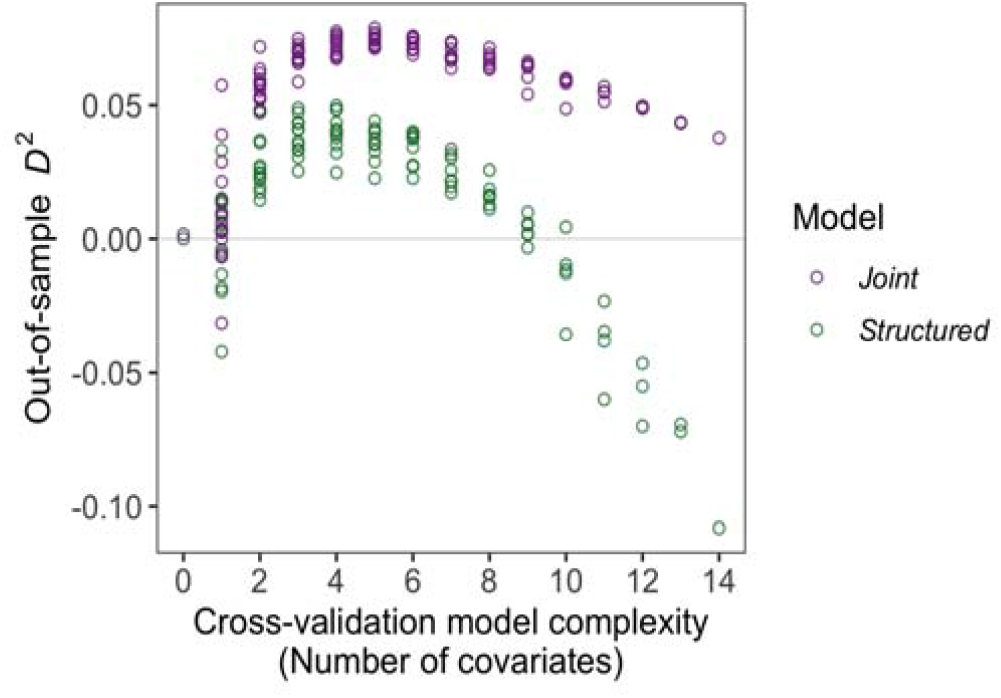
Out-of-sample during cross-validation with model complexity. The *Joint* model (purple) consistently outperformed the *Structured* model (green) in predicting out-of-sample local communities, and showed less susceptibility to over-fitting.

The optimal variables for both models included growing degree days at a regional scale, and local effects of relative soil temperature and vegetation cover (Fig. 4a). The optimal *Joint* model also included regional effects of forest cover and road length, while the optimal *Structured* model included local effects of canopy type. For many variables, species showed a variety of responses, resulting in no clear aggregate effect as the 95% Highest Posterior Density Interval (HPDIs) included zero (Fig. 4a, horizontal points and lines; Supplemental Fig. S1). However, the aggregate effect of growing degree days was positive in the *Joint* model, and the aggregate effect of squared growing degree days was negative in both models. For relative soil temperature, vegetation cover, and mixed canopy, nearly all species with non-zero HPDIs showed positive responses (Supplemental Fig. S2). The *Joint* model reduced uncertainty in species’ responses compared to the *Structured* model across regional covariates, as indicated by narrower 95% HPDIs (Fig. 4b). This reduction occurred on average across all species, but was greater for species not found in **Y**. In contrast, uncertainty in the *Joint* model was somewhat higher for local variables.

**Figure 4.**
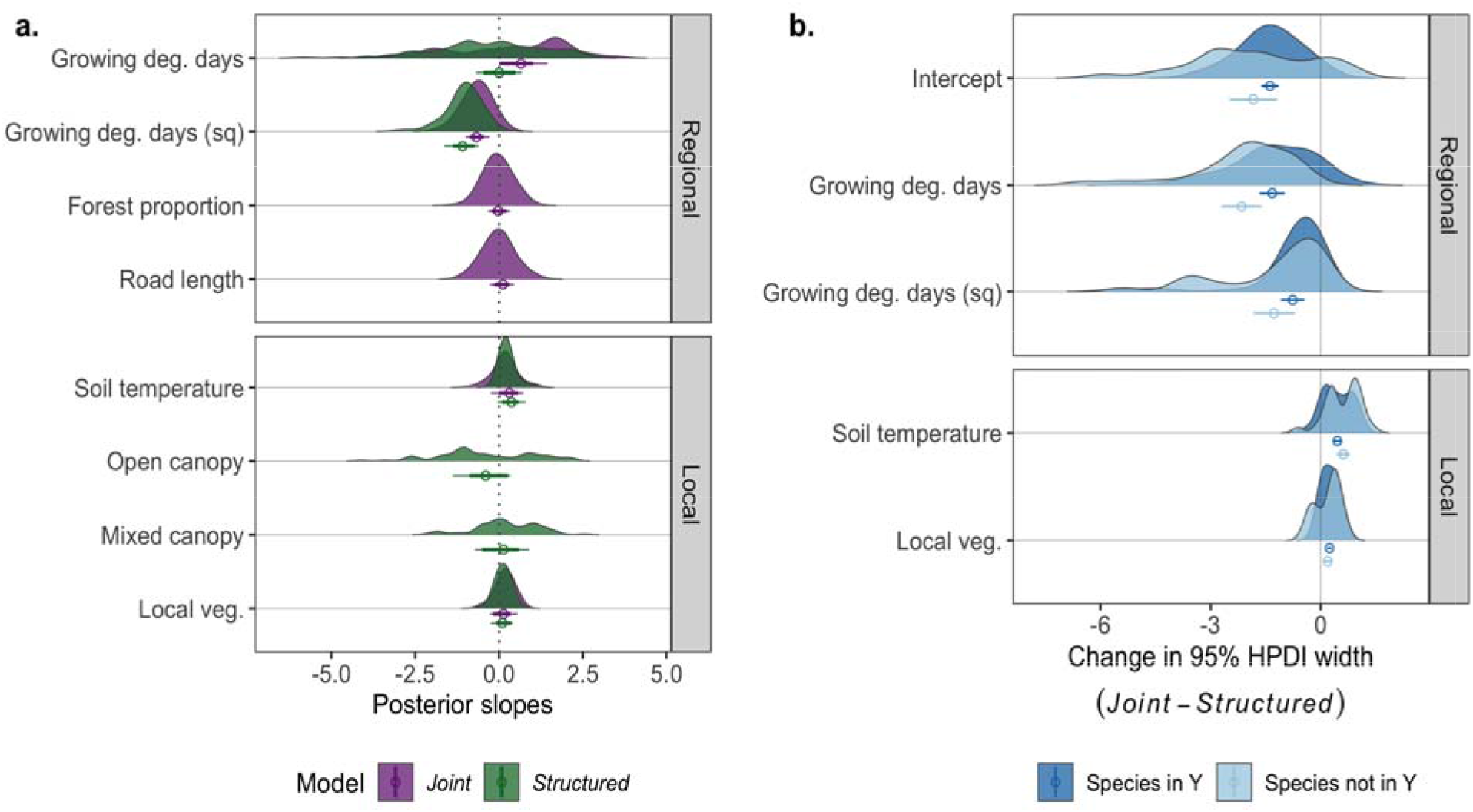
Covariate effects in optimal models. **(a)** Density curves summarise species-level posterior means for responses to variables in the model using both datasets (*Joint*: purple) and the model using only the abundance data (*Structured*: green). Points and lines show posterior means and Highest Posterior Density Intervals (HPDI; thick: 80%; thin: 95%) for the aggregate responses. **(b)** Change in 95% HPDI widths in species responses between models (negative: less uncertainty in *Joint* model) for species observed in the structured abundance dataset **Y** (dark blue) or only in the presence-only dataset **W** (light blue). Points and lines show mean standard error across species.

Both models predicted a peak in ant richness just below 1000m at a regional scale, with no clear trend at a local scale (Fig. 5a). In contrast, the *Joint* model predicted higher regional Shannon diversity at lower elevations, while the *Structured* model predicted little elevational trend (Fig. 5b). At a local scale, the *Structured* model predicted lower Shannon diversity at low elevations, increasing until 1000m, while the *Joint* model predicted little trend. The *Structured* model predicted higher regional colony intensities at lower elevations, in contrast to the increase in intensity above 1000m predicted by the *Joint* model (Fig. 5c). Local predicted intensity was similar across models, with little trend.

**Figure 5.**
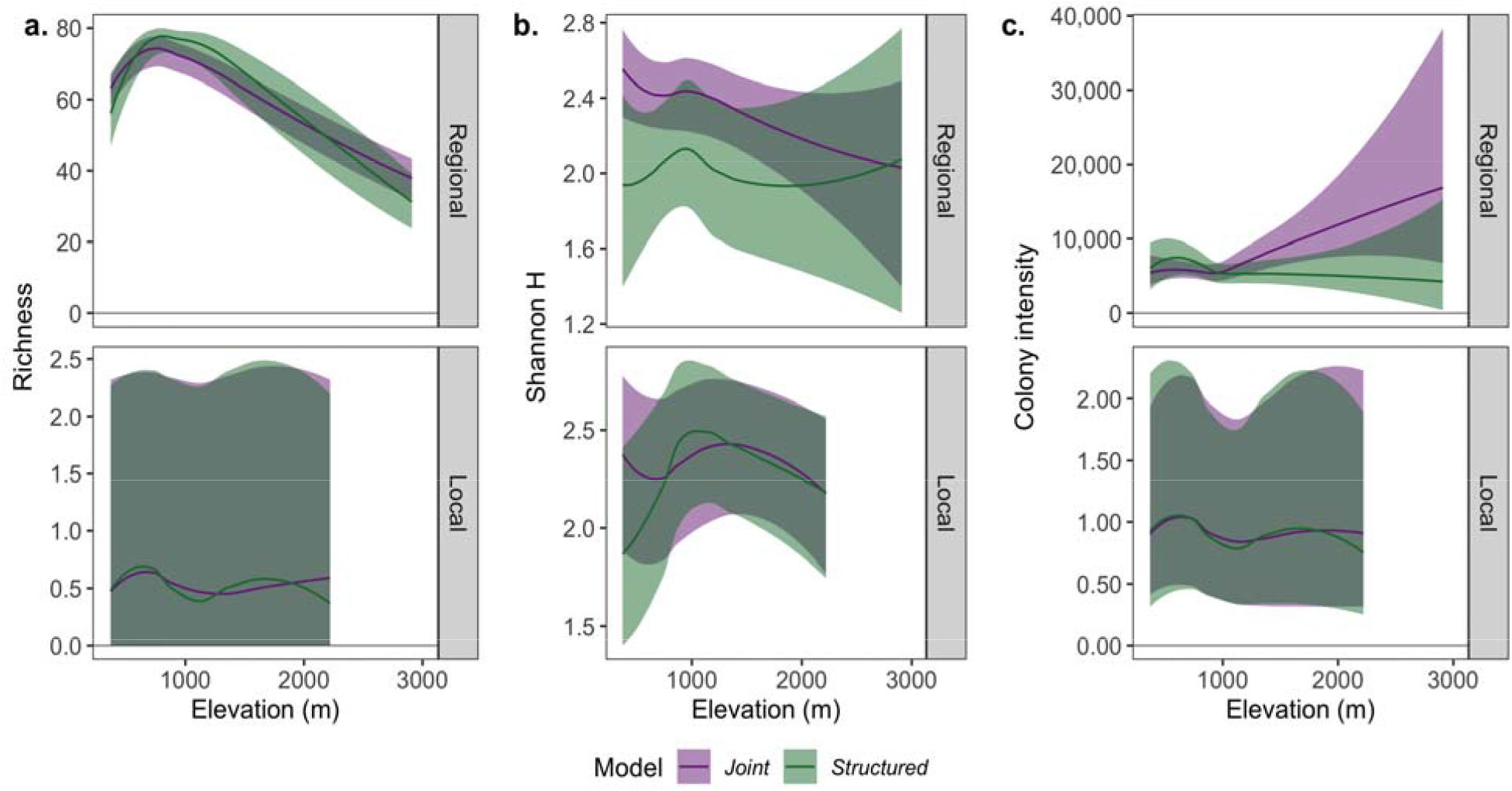
Elevational patterns of posterior distributions at regional and local scales for **(a)** species richness, **(b)** Shannon H diversity, and **(c)** total colony intensity. Lines and ribbons are loess lines using the posterior medians and 95% Highest Posterior Density Interval, respectively, for the model parameterized with both datasets (*Joint*: purple) and the model parameterized with only the structured abundance data (*Structured*: green).

Local ant communities showed strong separation across elevations. The DPCoA showed two distinct clusters representing plateau (1000m) and montane (1000m) plots, with results qualitatively similar for both models (*Joint*: Fig. 6, *Structured*: Supplemental Fig. S3). No clear pattern was seen at lower elevations, with broad, partially overlapping clusters from 300–700m and 700–1000m (Fig. 6a). Above 1000m, communities showed a gradient, with elevation increasing down the y-axis. Plateau communities formed a notably broad cluster (Fig. 6b), indicating greater variation among local ant communities. In contrast, montane communities clustered more tightly, with the western Jura mountain plots appearing largely as a subset of those in the taller, eastern Alps. Correspondingly, many species were predicted to occur predominantly in plateau or in montane zones (Supplemental Fig. S4), though 41 species (52%) exhibited elevational ranges that spanned both (Supplemental Table 3).

**Figure 6.**
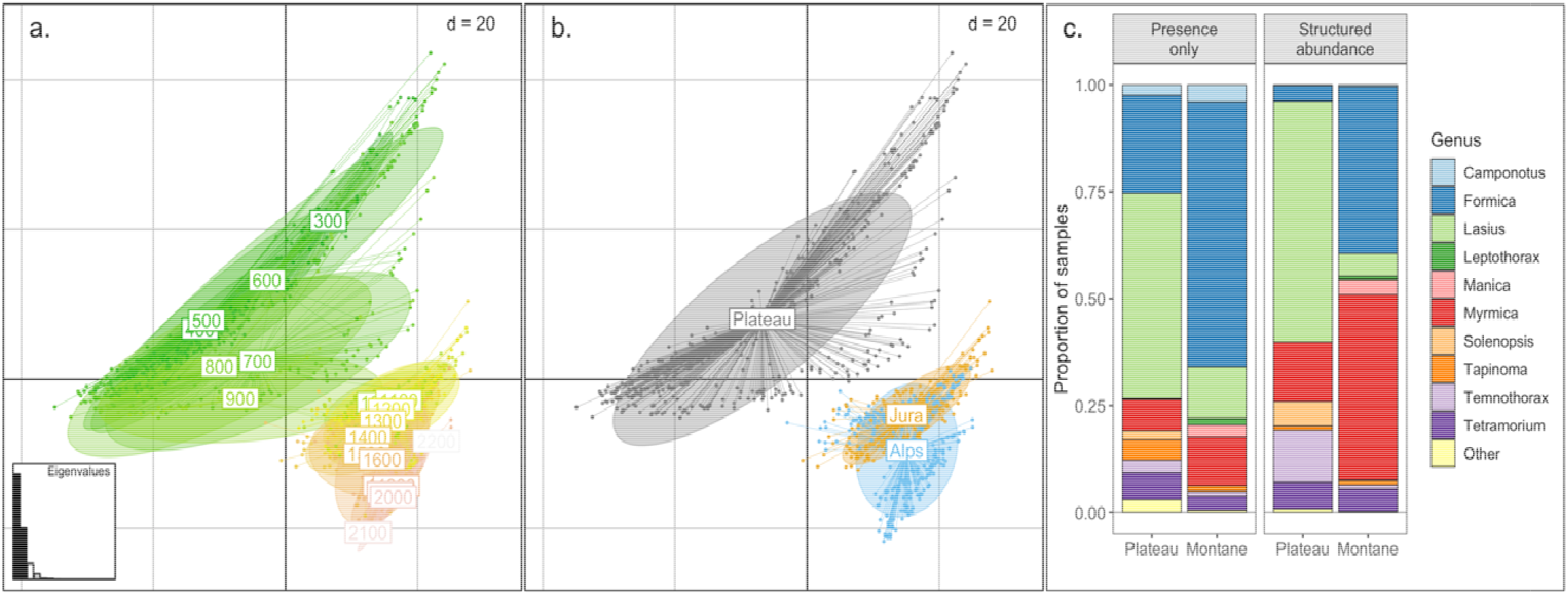
Elevational assemblages in the ant communities. Double principal coordinate analysi (DPCoA) of local communities predicted by the *Joint* model showed clustering by **(a)** elevational bin and **(b)** region. The central plateau extends to 1000m, with the Jura and Alp rising steeply in the west and east, respectively. **(c)** Genus composition in the presence-only and structured abundance datasets each showed differentiation across plateau and montane environments. Only genera that constitute ≥ 1% of at least one subset are shown, with others indicated as ‘Other’.

Both datasets showed strong differences in the genus composition between the plateau and montane samples (Fig. 6c). The plateau (<1000m) was dominated by *Lasius spp*., with high representation of *Formica spp*. in the montane zone (≥1000m). The structured abundance dataset **Y** also showed high relative abundance of *Myrmica spp*., particularly in montane environments. Compared to **Y**, the presence-only dataset **W** under-represented many species (95% HPDIs < 0; *Joint*: 23 species = 29%; Supplemental Fig. S5, including nearly every species in the genus *Myrmica*. In contrast, only the anthropophilic pavement ant *Tetramorium immigrans* was clearly over-represented (95% HPDIs > 0; *Joint*: 1 species = 1%; Supplemental Fig. S5).

Patterns of *β*-diversity among plots within structured sites did not dramatically differ between models (Fig. 7). Overall *β*-diversity showed weak elevational patterns (*Joint*: p=0.03, R^2^=0.10,*Δ*AICc=0.7; *Structured*: p=0.06, R^2^=0.06, *Δ*AICc=3.1), with relatively high variability among local communities on average regardless of elevation. At low elevations, the *balanced variation* and *abundance gradient* components were comparable, indicating differences in total intensity and in turnover of species identities among plots (*Joint*: p <0.001, R ^2^ =0.70, *Δ*AICc=5.4; *Structured*: p < 0.001, R^2^ = 0.48, *Δ* AICc=0.31). At higher elevations, the *abundance gradient* component dominated; plots differed in total intensity rather than in species identities (*Joint*: p < 0.001, R^2^=0.72,*Δ*AICc=10.6; *Structured*: p<0.001, R^2^=0.44, *Δ*AICc=2.4).

**Figure 7.**
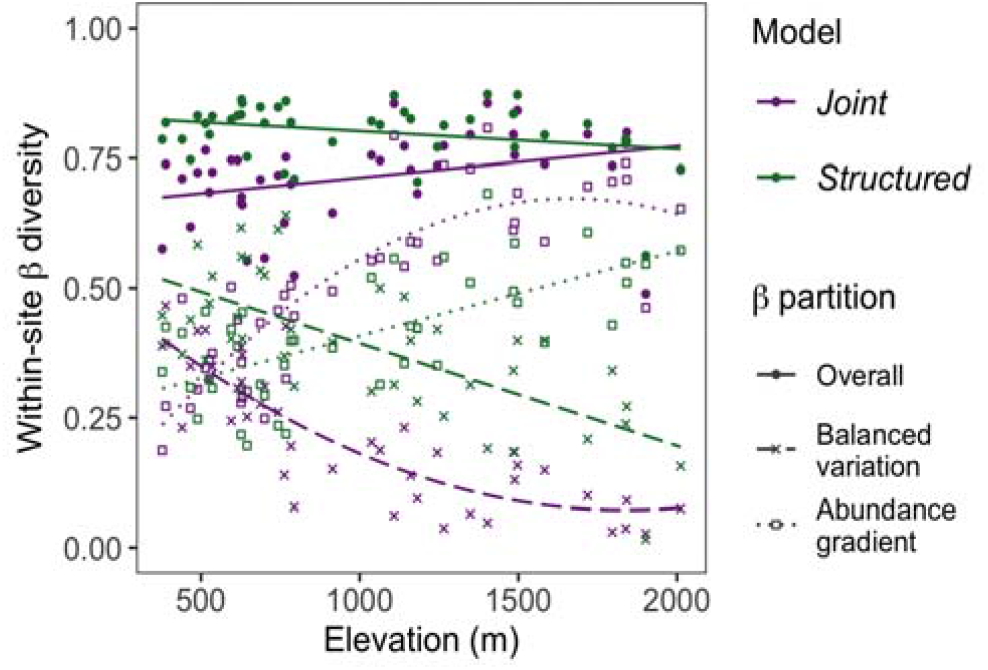
Multi-site -diversity and its components across elevation. Posterior medians were used to calculate multi-site -diversity and its components within each 1 km site, describing variation among plots. *Balanced variation* quantifies changes in relative abundance among species, while *abundance gradient* denotes changes in total abundance.

## Discussion

Samples collected by citizen scientists are part of many ecological datasets (Poisson et al. 2020). The *Joint* model integrating both datasets outperformed the *Structured* model based on the scientific sampling alone in predicting out-of-sample communities and was less susceptible to over-fitting. The information provided by the presence-only dataset at a regional scale produced more accurate, robust predictions at a local scale. Further, the *Joint* model reduced uncertainty in species-level regional responses, most dramatically for species that were not detected in the structured abundance dataset. In the *Joint* model, regional ant richness was predicted to peak at the base of the mountains, declining in both lower and higher elevations. In contrast, Shannon diversity was predicted to decline somewhat from low to high elevations. Patterns were best predicted by growing degree days, forest cover, and road length at a regional scale, and soil temperature, understory vegetation cover, and canopy cover at a local scale. Local ant communities showed plateau and montane clusters, with distinct changes in the generic composition between zones.

Ant richness frequently peaks at intermediate elevations, as observed here and in other Alpine regions (Hellrigl 2003; Glaser 2006; Szewczyk and McCain 2016). While average local plot abundance, richness, and diversity remain largely stable across the gradient, the regional scale decrease in richness and increase in colony intensity at high elevations implies density compensation (Longino and Colwell 2011). However, there was great uncertainty in predicted regional intensity at high elevations, especially in the *Joint* model. Several factors likely contribute. The presence-only dataset included more observations of mound-building montane species that can occur at extremely high densities, such as *Formica pressilabris* (Seifert 2018). Additionally, intensity magnitudes are informed by the local plots, which only extended to 2222 m, while the regional intensities were predicted as high as 2912 m. These elevations contain novel conditions beyond those that exist at the structured abundance plots, and so the fitted relationships are extrapolated. For example, several mound-building montane species of *Formica* showed strong negative responses to roads (Supplemental Fig. S1), which are sparse at high elevations. The large uncertainty in the *Joint* model predictions thus reflect the interplay of novel physical environments, limitations of the local abundance data, and species’ ecology (Harris, Taylor, and White 2018; Guillera-Arroita, Kéry, and Lahoz-Monfort 2019).

At both regional and local scales, temperature is a key determinant of the structure of the ant community, consistent with much past research (Sanders et al. 2007; Penick et al. 2017; Bishop et al. 2017; Szewczyk and McCain 2016; Szewczyk and McCain 2019). However, thermal preferences were highly variable between species, including among congeners (Supplemental Fig. S1). Responses to canopy, forest cover, and road length were similarly variable, indicating differentiation of broad habitat type preferences among species. In contrast, species with clear responses to soil temperature or local vegetation coverage tended to occur in relatively warm and relatively productive microhabitats (Supplemental Fig. S2). The ant species most affected by local productivity preferred denser vegetation (Andersen 1986; Kaspari, O’Donnell, and Kercher 2000). Nevertheless, only select taxa showed positive relationships, with no effect on the predicted colony intensity of most species. Specifically, many *Myrmica* species, *Solenopsis fugax*, and two *Lasius* species (*L. niger, L. flavus*) were all more likely to occur in more productive microhabitats. Similarly, a relatively small number of species showed strong positive or negative effects of canopy cover.

Plateau (<1000 m) and montane (> 1000 m) local communities were quite distinguishable based on species composition, and tended to form clear clusters (Fig. 6). However, the communities do not appear to be strongly Clementsian such that there is little overlap between them; instead, 52% of species were detected in both the plateau and the montane, with only 33% restricted to the plateau and 15% exclusively montane, and the observed and modelled elevational ranges show many independently distributed species rather than cohesive plateau and montane assemblages (Supplemental Table 3, Supplemental Fig. S3). Local communities also showed more variation in species composition at lower elevations, with greater turnover in species identities among plots. While total *β*-diversity was comparable at high elevations, the difference among plots was driven by changes in the total number of expected colonies rather than by which species were likely to occur.

The model described here, like any, required assumptions. The predicted intensity at 1 km^2^ assumes that the plots within each structured abundance site are representative of the larger area. Spatially discrete models require summarising the variation within each spatial unit, and consequently local variation is not accounted for (Calabrese et al. 2014; Frishkoff, Mahler, and Fortin 2019; Amini Tehrani, Naimi, and Jaboyedoff 2020). While the plots are reasonably representative within the 44 structured sites (Fig. 1), extrapolation to the larger grid may fail to account for more extreme heterogeneity in suitability, such that the modelled regional intensity and richness may be better interpreted as an upper bound (Guillera-Arroita, Kéry, and Lahoz-Monfort 2019; Altwegg and Nichols 2019; Johnston et al. 2020). In fact, past work suggests that an underestimation of heterogeneity should be expected, both in individual species’ distributions and in the overall patterns of richness, abundance, and diversity, resulting from error in the estimation of species’ responses, error in the included predictors, or predictors that were not included (Calabrese et al. 2014; Biber et al. 2019; Guillera-Arroita, Kéry, and Lahoz-Monfort 2019). Though the local latent variable attempted to model some combination of missing predictors or biotic interactions, an unfeasibly large number of latent variables may be necessary to accurately capture local dynamics (Ovaskainen et al. 2016).

A key attribute of jointly estimated models is that information can be shared across species. This technique, known as hierarchical shrinkage, leads to improved predictions, particularly for species with sparse data (Fithian et al. 2015). Here, species with little or no data were assumed to show preferences similar to congeners. Thus, in the *Structured* model, species undetected in **Y** were predicted to be rare, but to show similar relative responses as other species within the same genus. For example, taxa that do not nest in the soil (e.g., *Camponotus spp*.) were very unlikely to be detected in the structured soil plots (Supplemental Table 3). Consequently, the *Structured* model predicted that *Camponotus* species were generally rare with a relative pattern informed by the aggregate (*β*) responses as well as the limited detections of *C. ligniperdus* (Supplemental Figs. S2, S6). In contrast, the *Joint* model predictions were rather different, because the preferences of these species were further informed by **W** (Supplemental Figs. S1, S7). Nevertheless, species with few total observations were more similar to the genus-level average. Thus, given a paucity of species-level data, hierarchical shrinkage provides a ’best guess’ based on other species, with correspondingly large uncertainty (Fig. 4b, Supplemental Fig. S1).

The presence-only dataset **W** showed clear spatial and taxonomic bias. These are common issues in citizen science projects (Ward 2014; Theobald et al. 2015; Johnston et al. 2020; Robinson et al. 2020), and the model structure explicitly incorporated these biases (Isaac et al. 2014). Though coverage was expansive through the region (Fig. 1), many grid cells were represented by a single sample, with a few cells containing more than one hundred. Similarly, the taxonomic composition of **W** was not fully aligned with **Y**, which more accurately quantified colony density (Supplemental Fig. S5, Supplemental Table 3). While some anthropophilic species were likely over-represented, **W** showed a stronger tendency toward under-representation. Many of the under-represented taxa have somewhat unobstrusive lifestyles or tend to live in less trafficked habitats (Seifert 2018), nesting and foraging in denser vegetation (e.g., some *Myrmica* species), under the soil (e.g., some *Lasius* species), or in twigs (e.g., *Temnothorax* species). Further, these taxa are not so rare or morphologically unique that they would be specifically sought after by contributing experts or enthusiasts.

Despite these biases, the presence-only dataset also made clear contributions. The citizen science efforts greatly expanded the taxonomic breadth of the survey, including 28 species (35% of total) and 7 genera (33% of total) that were not detected in **Y** (Supplemental Table 3). Opportunistic datasets often better capture uncommon species (Henckel et al. 2020). Correspondingly, the taxa detected only in **W** tended to be rare, including 9 singletons and 4 doubletons. The integration of the two datasets consequently allowed for more precise estimation of species responses (Fig. 4b), particularly for species with few observations. As a result, the presences, incorporated into the model only at a regional scale, improved out-of-sample predictions at a local scale (Fig. 3). The presence-only dataset also rendered the *Joint* model less susceptible to over-fitting. In contrast, the *Structured* model declined in predictive performance at rather modest levels of complexity, performing worse than the intercept-only model when more than 9 covariates were included, while the *Joint* model showed only limited declines in performance.

The efforts of citizen scientists in the canton of Vaud thus contributed valuable information not captured by the concurrent structured samples. The broad spatial coverage captured a similarly broad array of species, including rarer species that were unlikely to be detected in a structured survey focused on colony density. The *Joint* model accounted for spatial and taxonomic bias, as well as geo-locational imprecision, to improve the estimation of species’ responses to environmental variables as well as the predicted distribution of ants across the region. By leveraging the strengths of each collection effort, the hierarchical model described here provides a framework for leveraging the efforts of citizen scientists to improve our understanding of the distribution of biodiversity.

## Acknowledgments

We offer thanks to all the citizen scientists who contributed to Opération Fourmis, as well as the researchers, staff, and expert taxonomists associated with the project. We further thank all the volunteers and technicians who dedicated hours to collecting the structured samples. Funding was provided by the Herbette foundation of the University of Lausanne, part of a bequest of M. Rullens to the University of Lausanne, the Société Vaudoise des Sciences Naturelles and the “Retraites Populaires”.

## Data and code availability

The code used for genetic species identification and the COI sequences are available at https://github.com/glavanc1/ants_ID. The code used for all other analyses is available at https://github.com/Sz-Tim/opfo and https://github.com/Sz-Tim/5_citsci

## Appendix Descriptions

### Supplemental methods

**Supplemental Table 1**. Land cover types and canopy classification

**Supplemental Table 2**. Prior distributions

**Code A.1**. Stan code for full model

### Supplemental results

**Supplemental Table 3**. Species list with observed elevational range and counts

**Supplemental Table 4**. Variable selection results

**Supplemental Figure S1**. Species-level responses (points + HPDIs)

**Supplemental Figure S2**. Species-level directional response summaries (bar proportions)

**Supplemental Figure S3**. Plateau vs montane species’ intensities

**Supplemental Figure S4**. DPCoA based on *Structured* model predictions

**Supplemental Figure S5**. Taxonomic bias ->S5

**Supplemental Figure S6**. Maps of predicted richness by genus (*Structured*)

**Supplemental Figure S7**. Maps of predicted richness by genus (*Joint*)

#### Supplemental Methods

**Supplemental Table 1:** Land cover types and canopy classifications with the number of plots, colonies, and species detected in the structured abundance dataset. The percent of the column total is shown in parentheses.

| Land cover type | Canopy | Plots | Colonies | Species |
| --- | --- | --- | --- | --- |
| Mixed forest | Closed | 82 (7.7%) | 68 (6.2%) | 14 (27.5%) |
| Deciduous forest | Closed | 97 (9.2%) | 68 (6.2%) | 15 (29.4%) |
| Conifer forest | Closed | 99 (9.3%) | 55 (5.0%) | 13 (25.5%) |
| Alluvial zone | Mixed | 3 (0.3%) | 4 (0.4%) | 2 (3.9%) |
| Orchards & vineyards | Mixed | 16 (1.5%) | 29 (2.7%) | 6 (11.8%) |
| Roadside | Mixed | 46 (4.3%) | 67 (6.1%) | 16 (31.4%) |
| Transition | Mixed | 71 (6.7%) | 70 (6.4%) | 23 (45.1%) |
| Quarry | Open | 2 (0.2%) | 1 (0.1%) | 1 (2.0%) |
| Gravel | Open | 4 (0.4%) | 4 (0.4%) | 2 (3.9%) |
| Marsh | Open | 6 (0.6%) | 11 (1.0%) | 5 (9.8%) |
| Wetland | Open | 7 (0.7%) | 3 (0.3%) | 3 (5.9%) |
| Scree | Open | 23 (2.2%) | 8 (0.7%) | 6 (11.8%) |
| Dry meadow | Open | 38 (3.6%) | 63 (5.8%) | 19 (37.3%) |
| Built zone | Open | 52 (4.9%) | 57 (5.2%) | 13 (25.5%) |
| Other open | Open | 513 (48.4%) | 582 (53.4%) | 35 (68.6%) |

**Supplemental Table 2:**
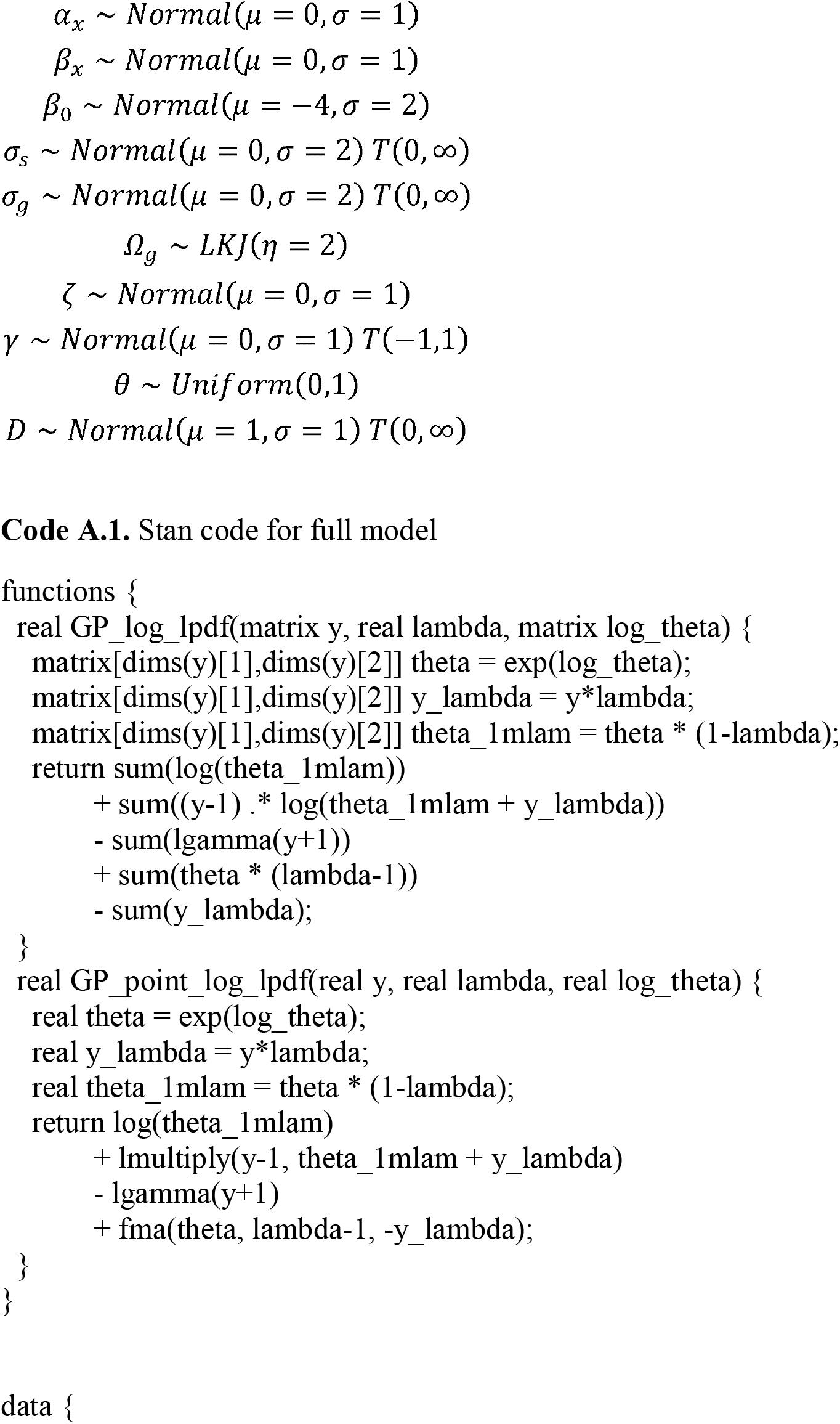
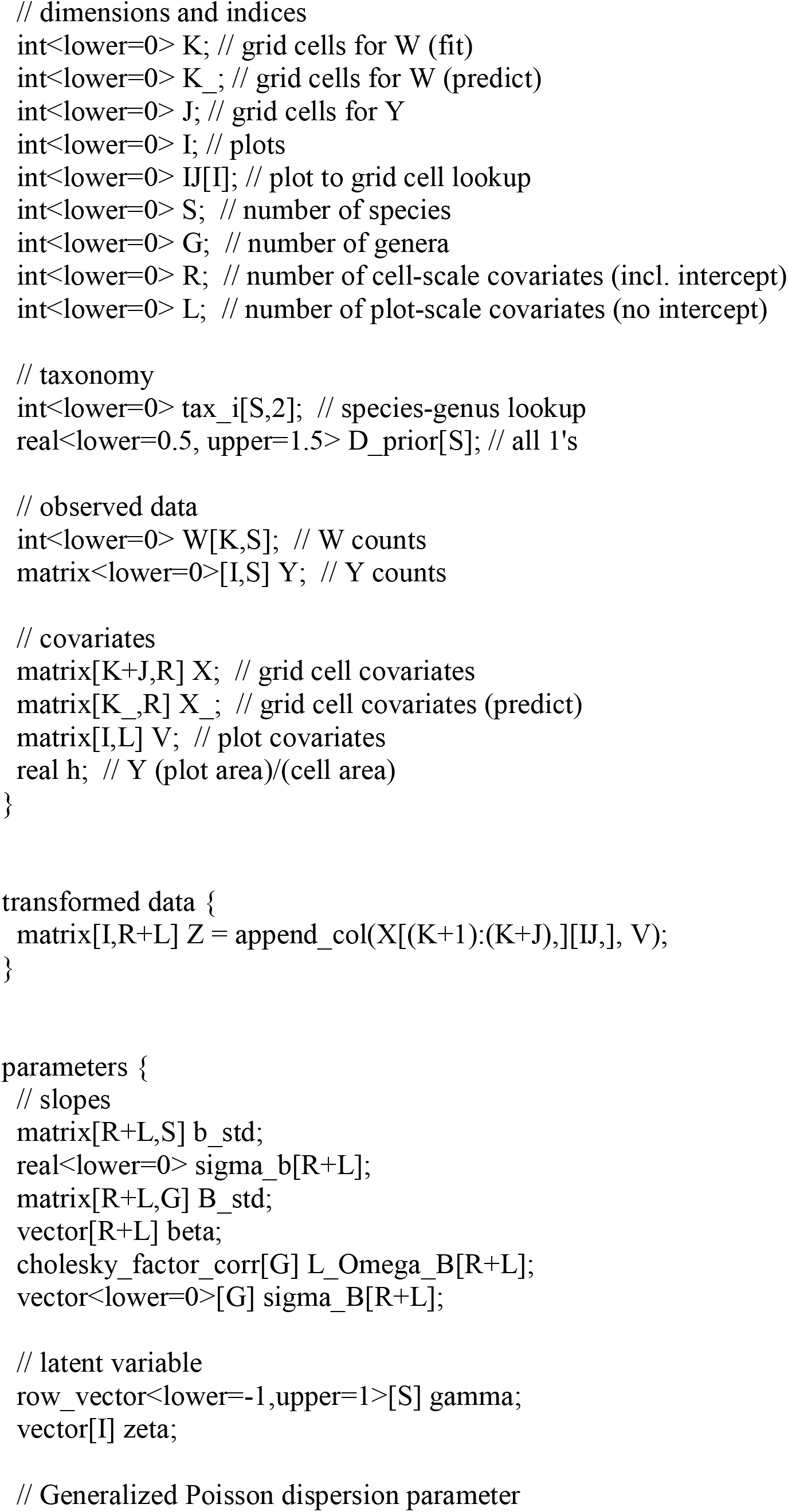
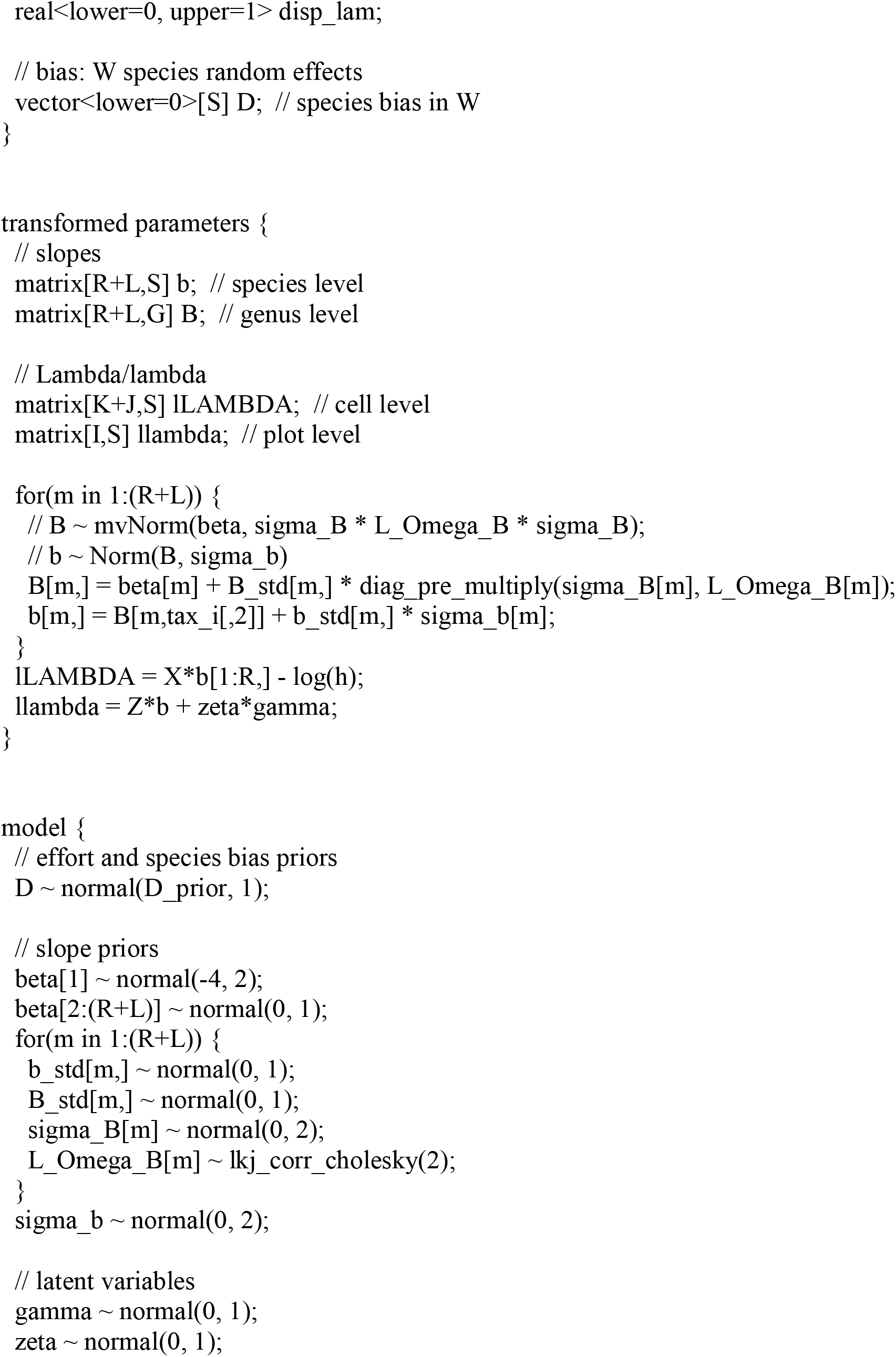
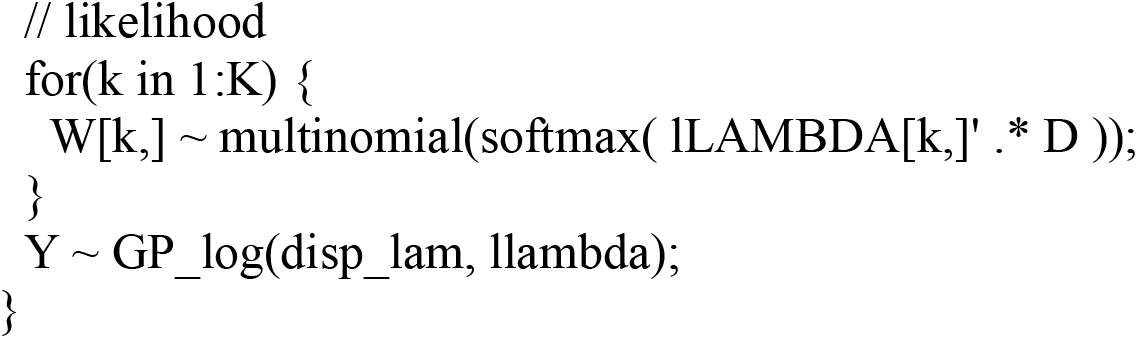
Prior distributions. Prior distributions were lightly informative, constraining values to plausible ranges to improve sampling. The correlation matrices for the genus-level slopes used a Cholesky decomposition and followed an LKJ prior as recommended for better model performance in Stan (Carpenter et al. 2017; Caradima, Schuwirth, and Reichert 2019).

### Supplemental Results

**Supplemental Table 3:**
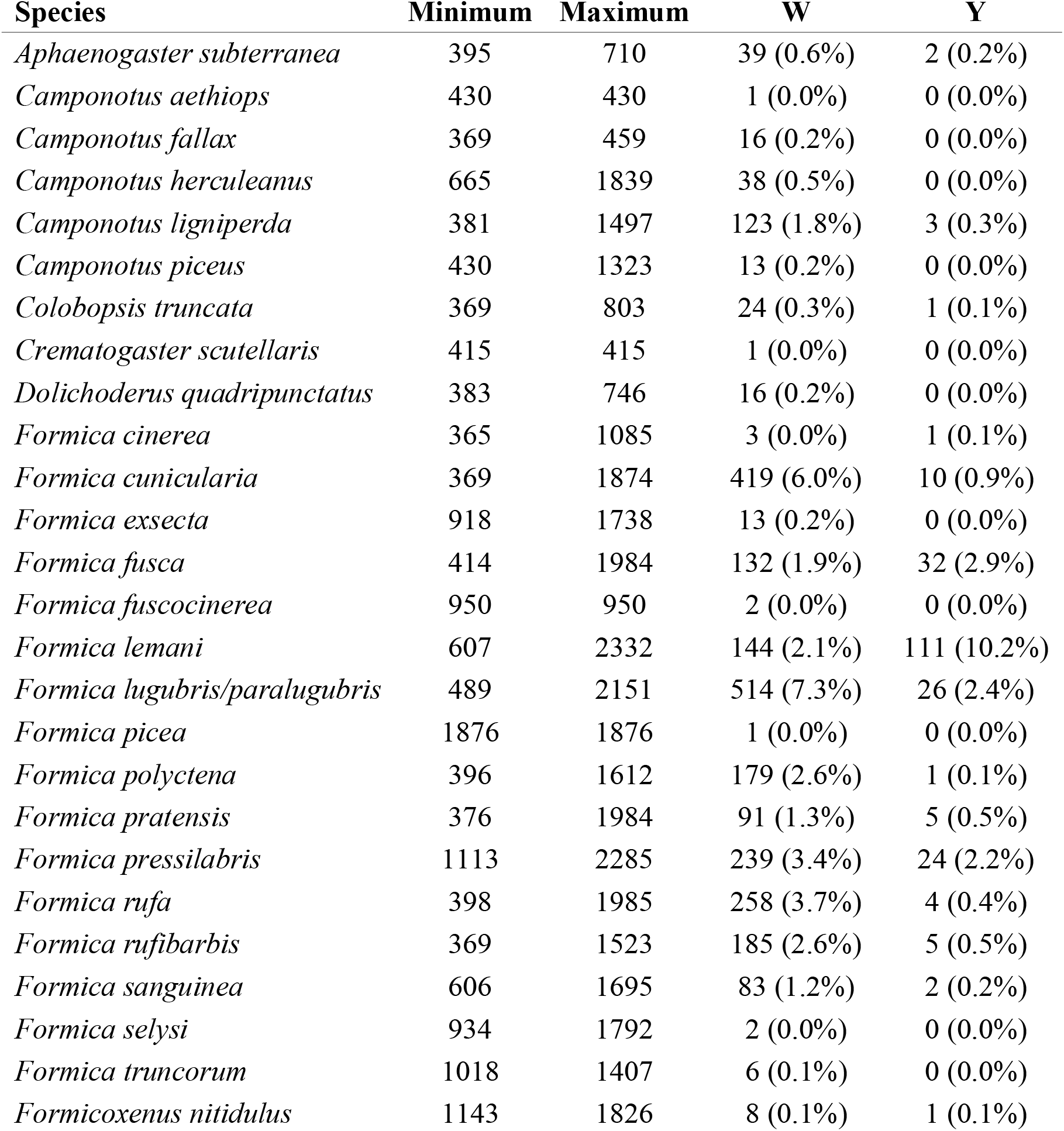
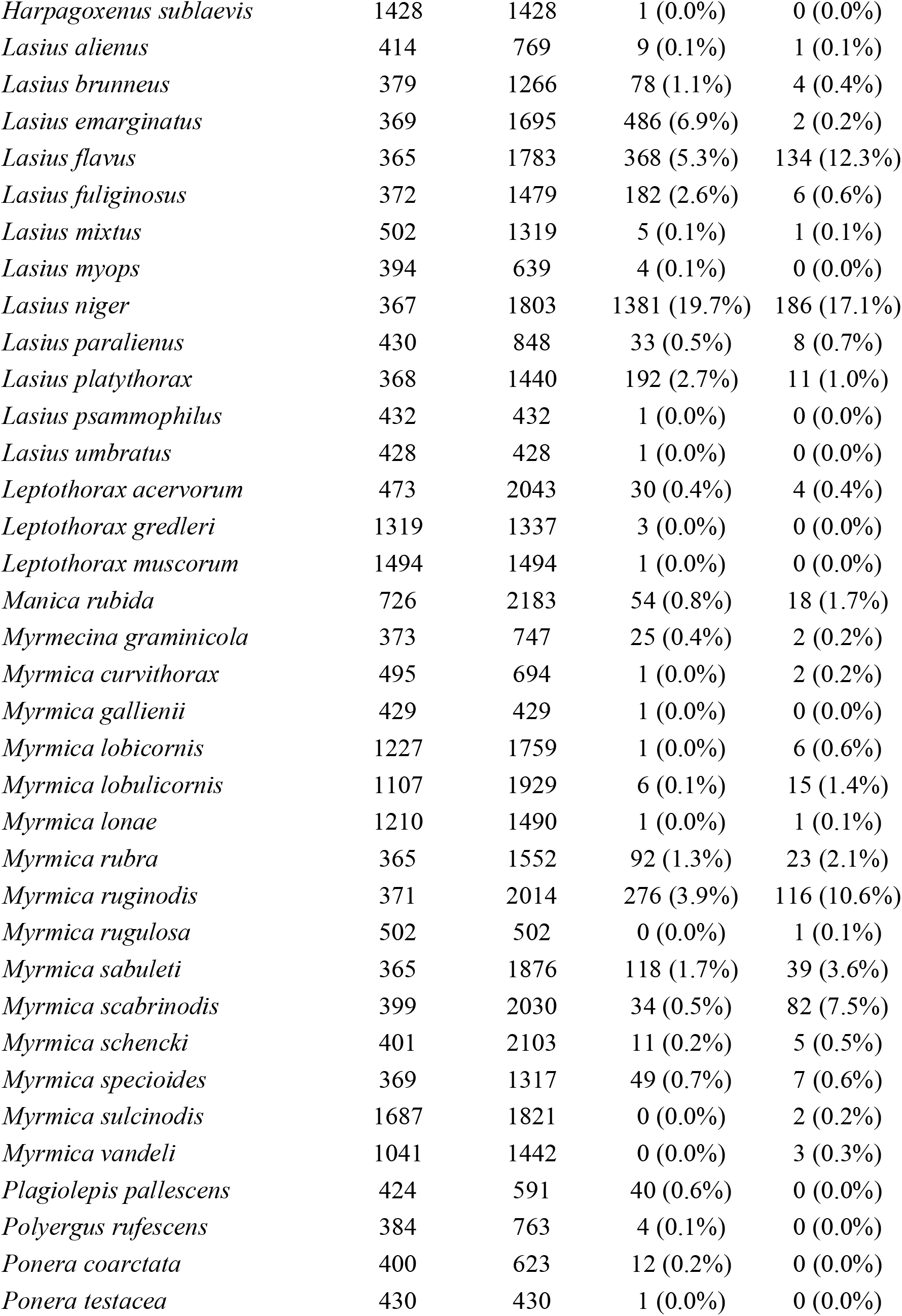
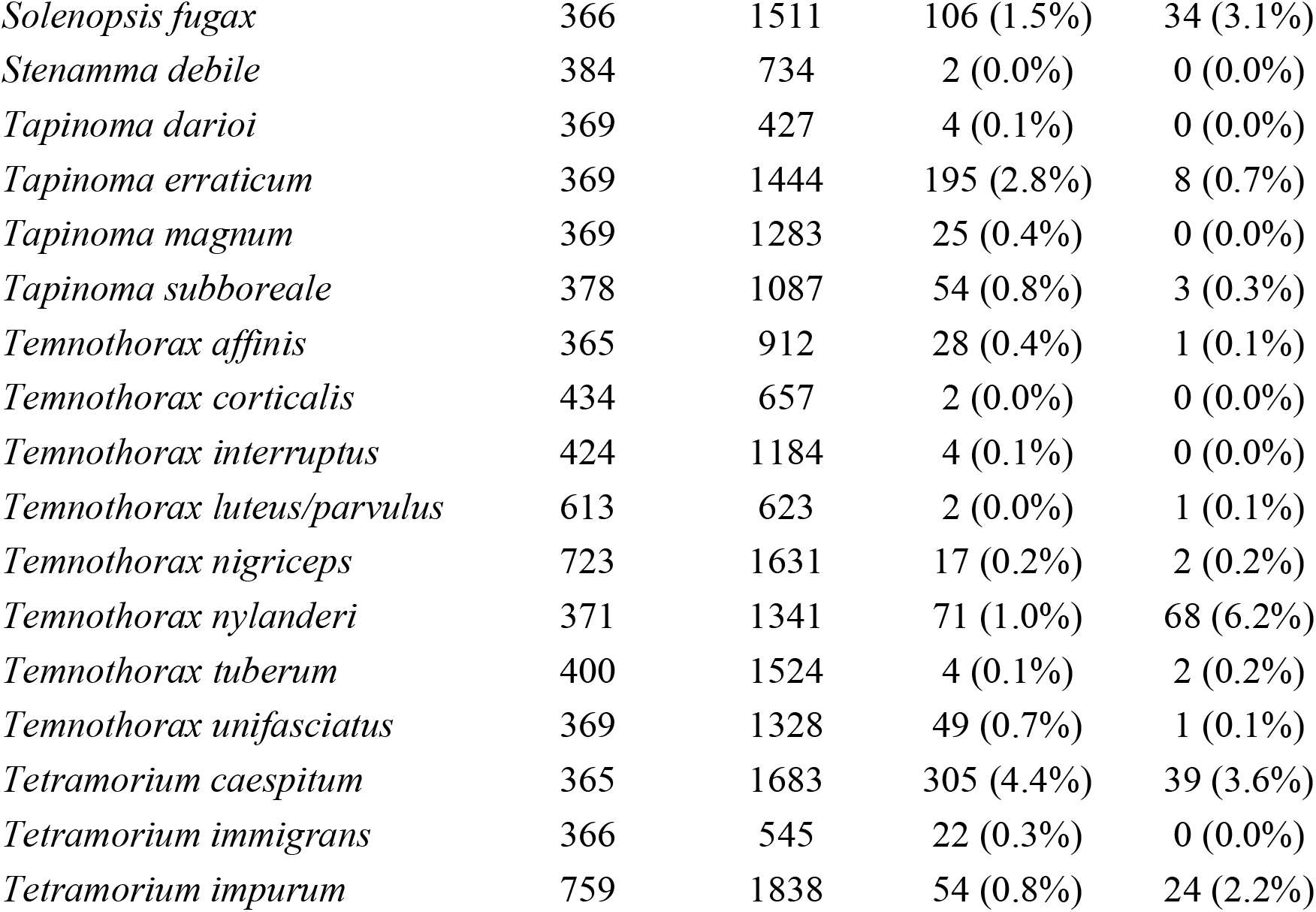

**Supplemental Table 4:**
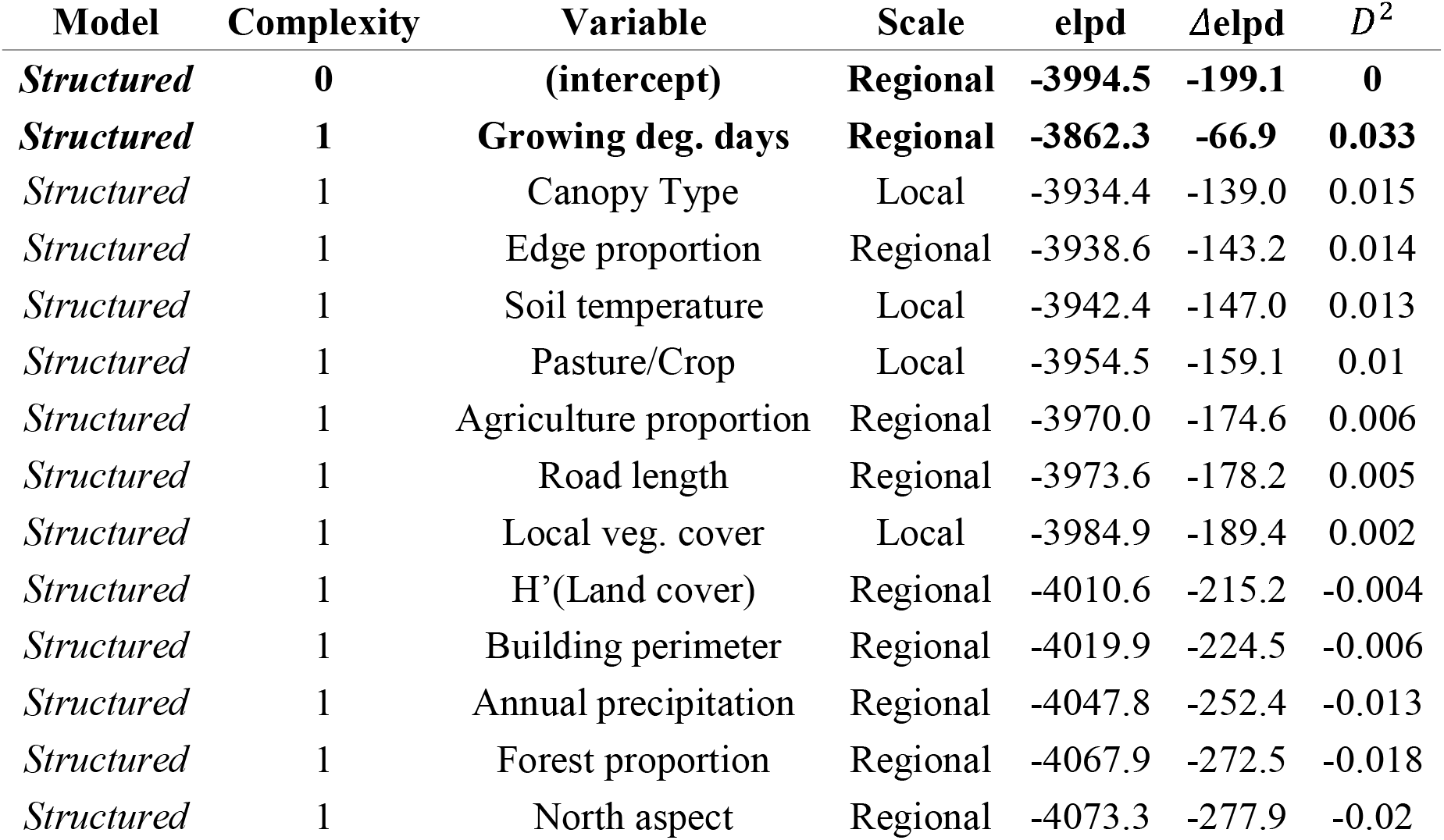
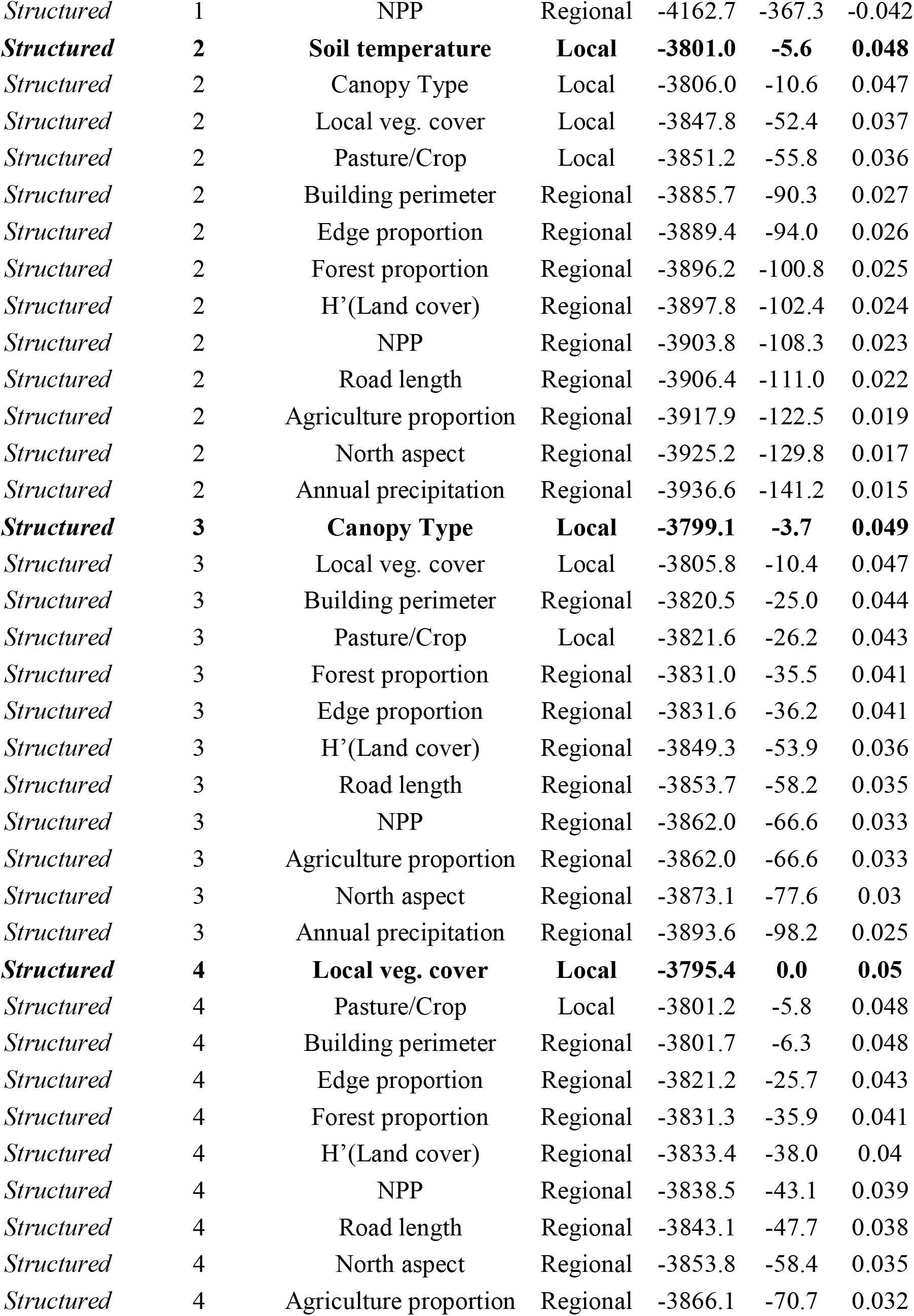
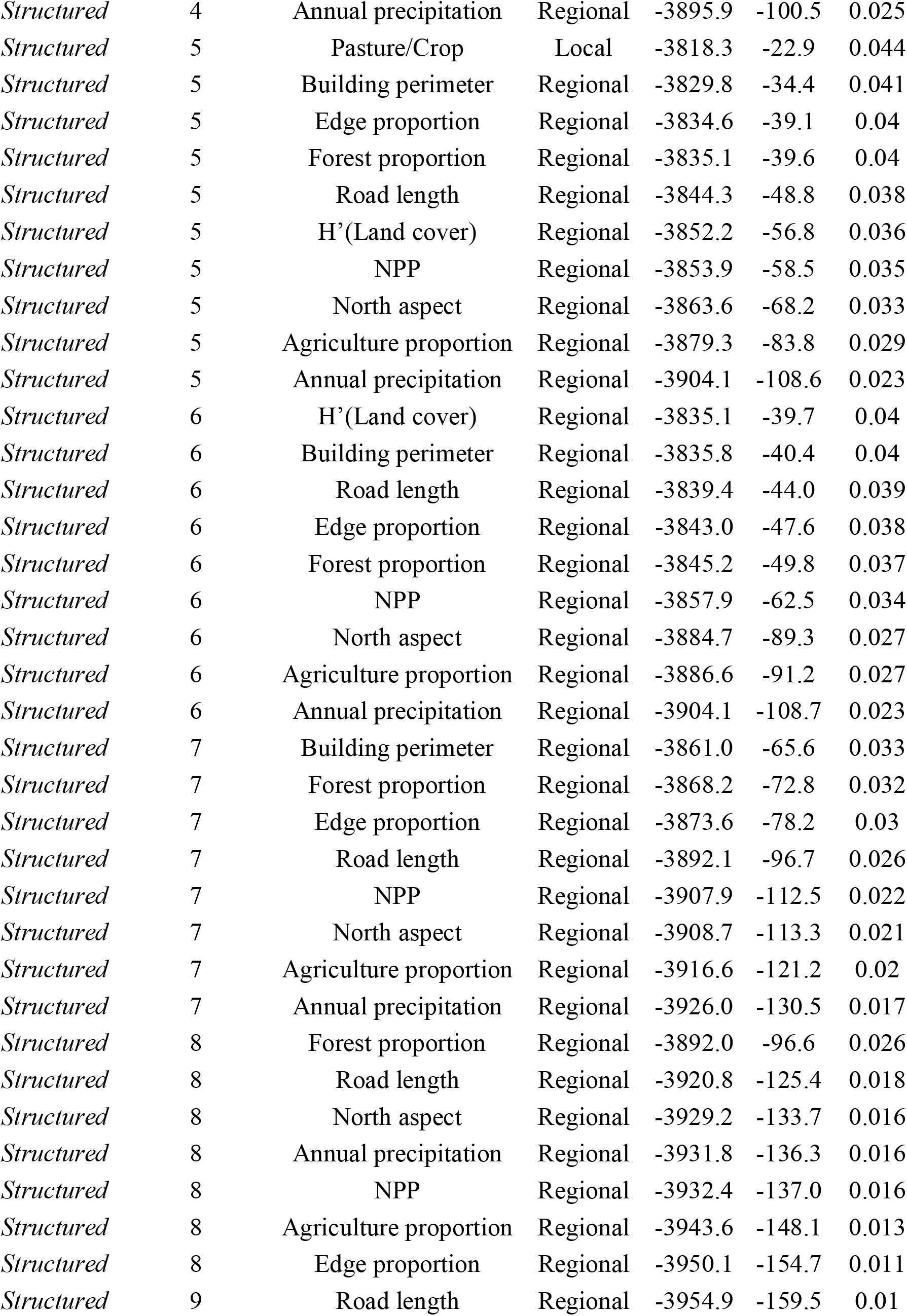
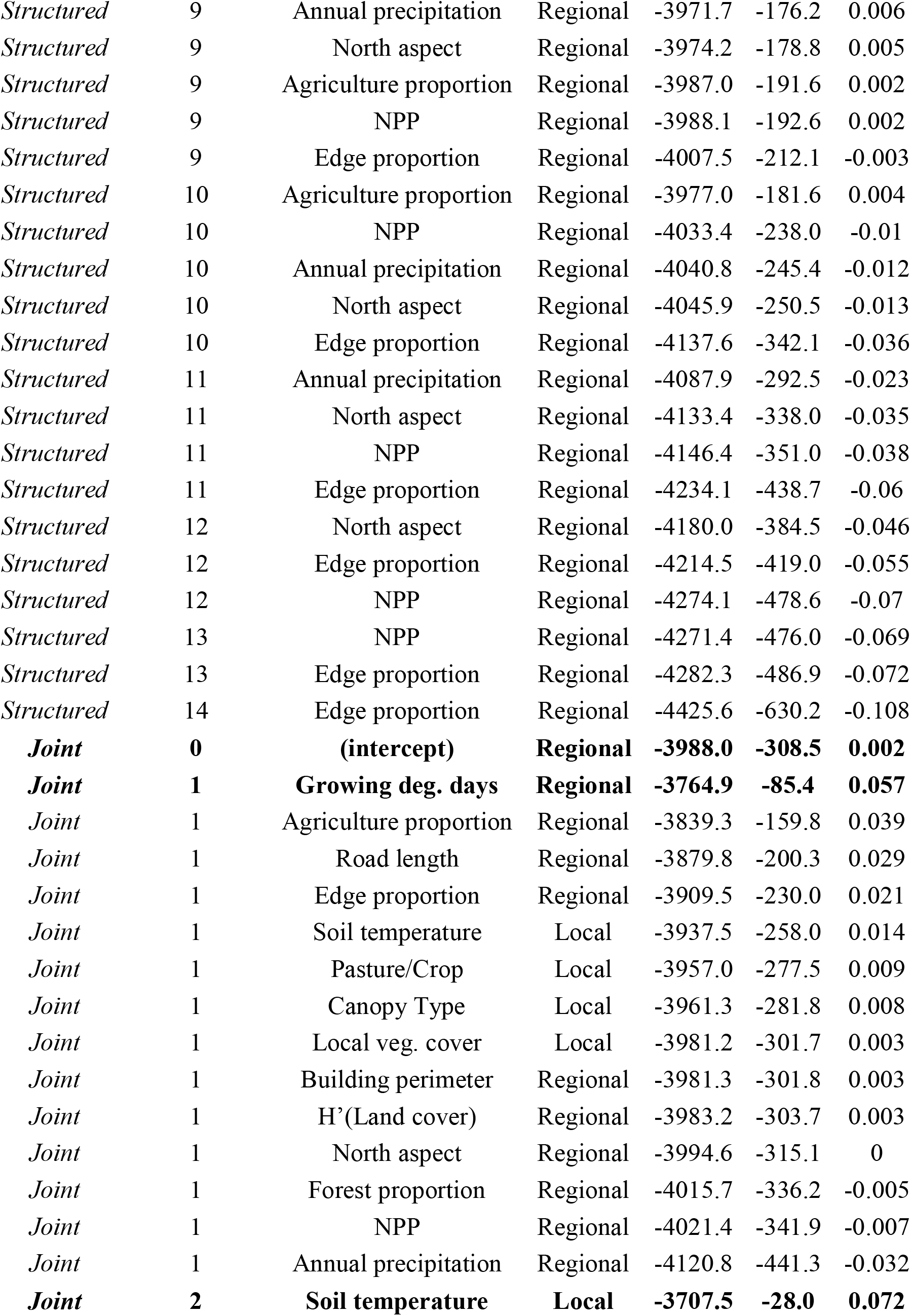
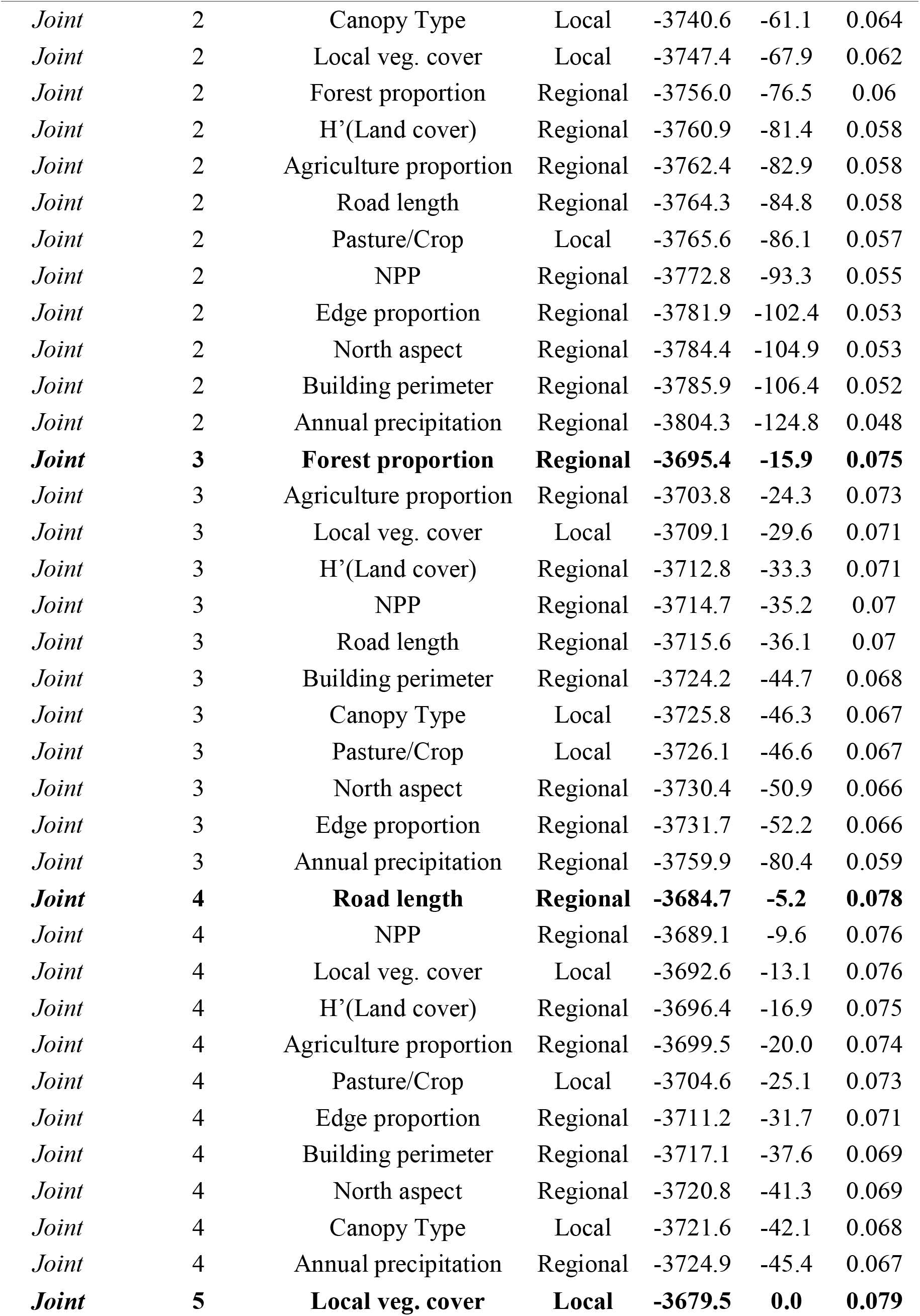
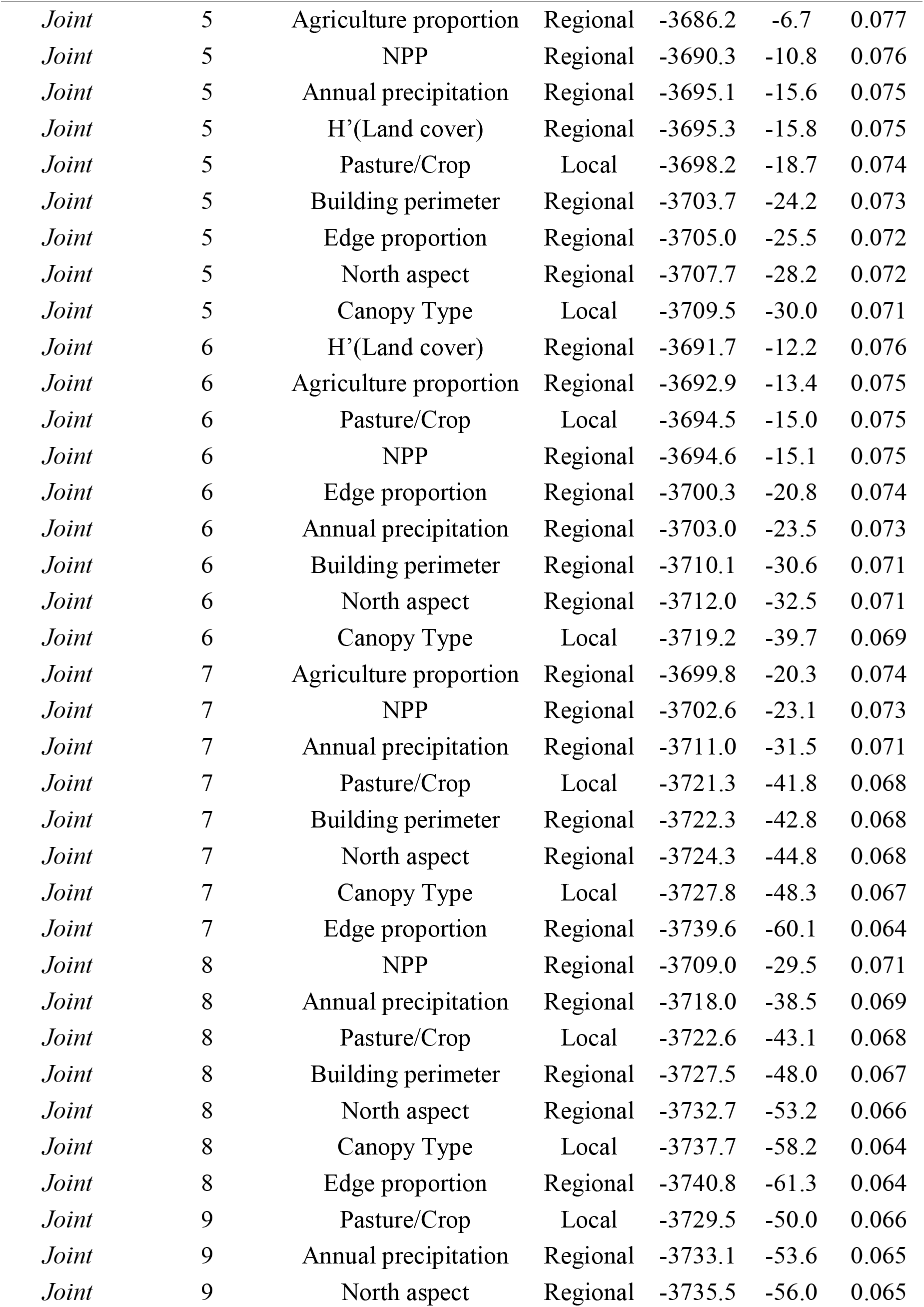
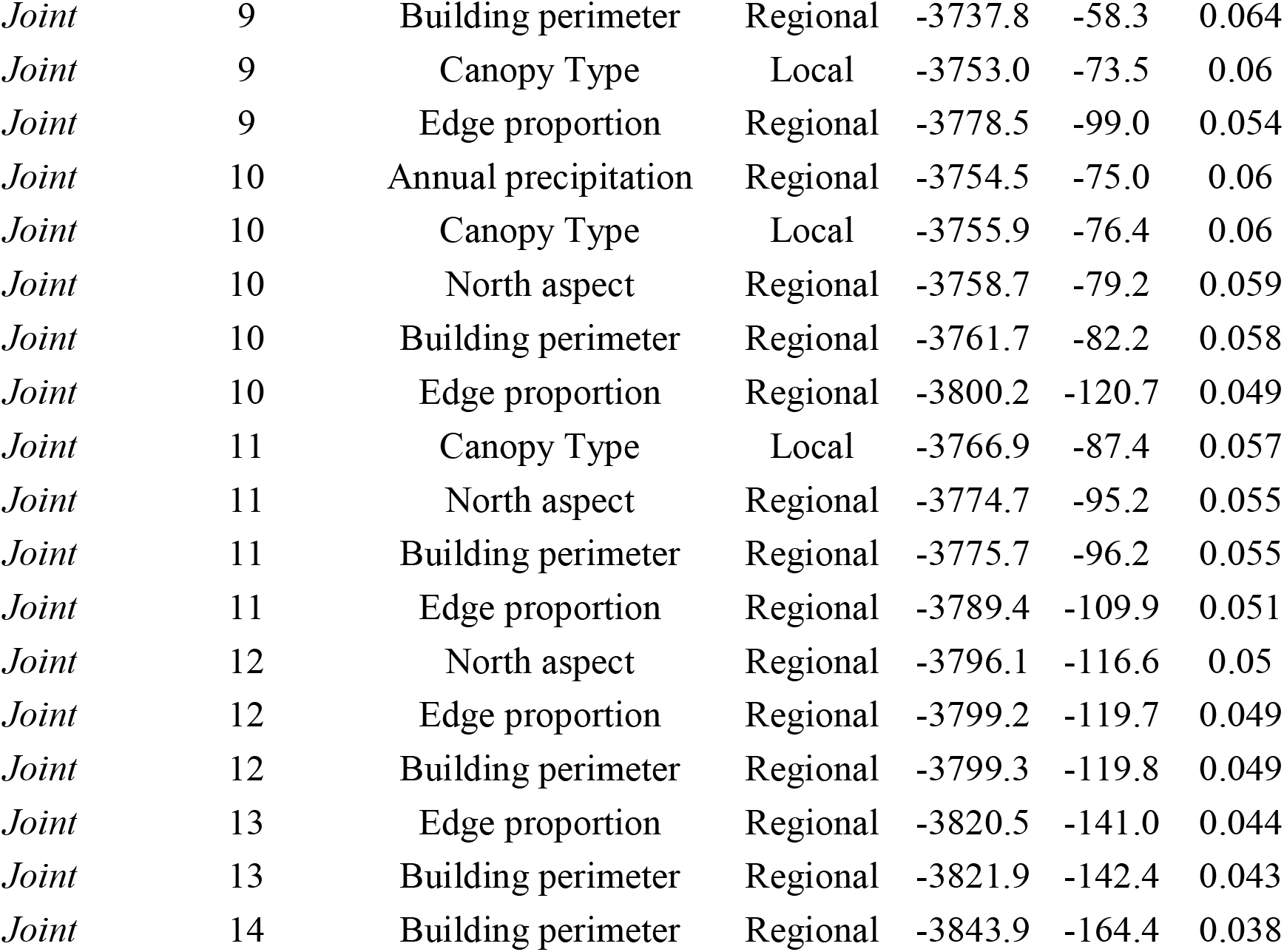

**Supplemental Figure S1:**
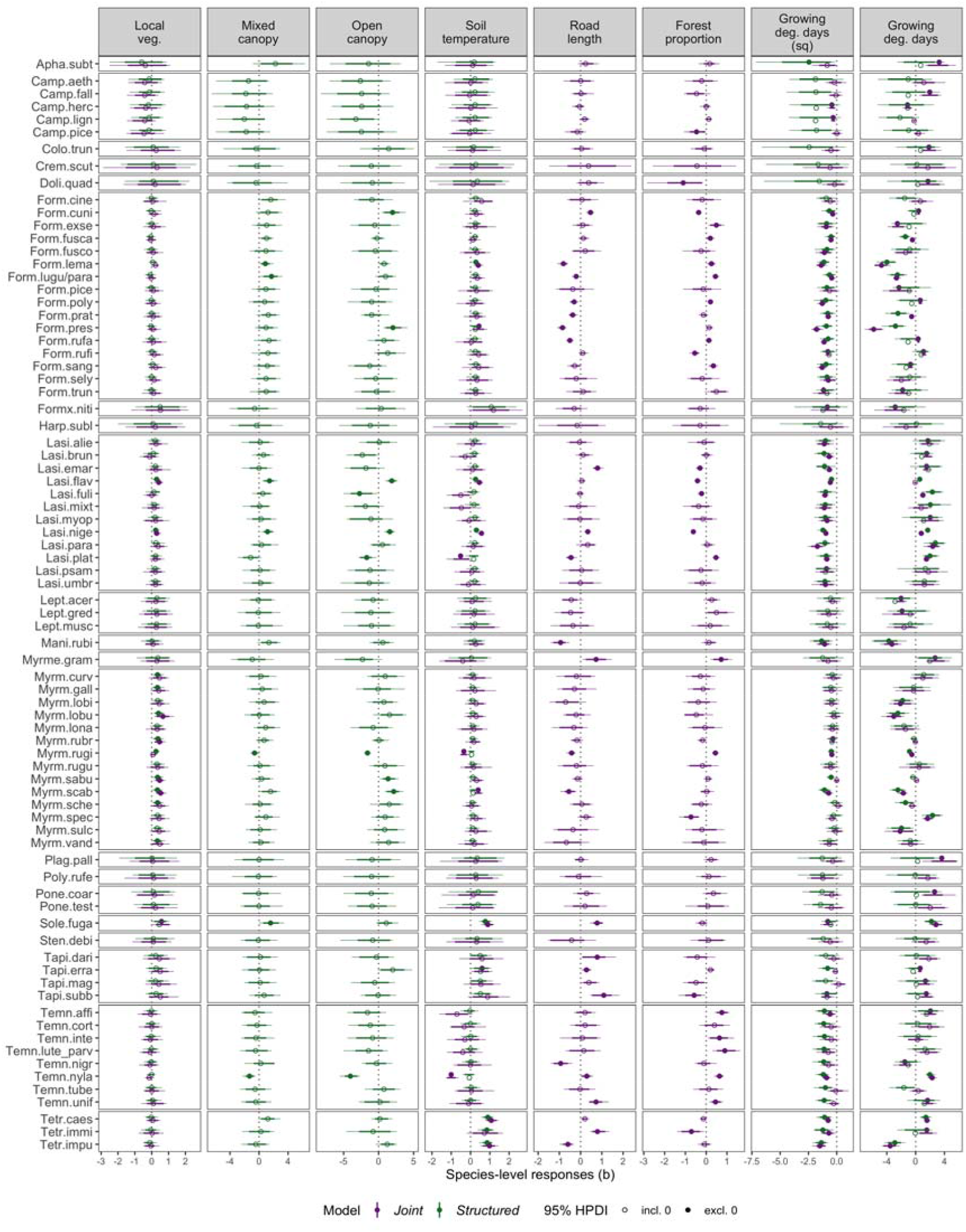
Posterior species responses to local and regional covariates. Points show posterior medians, while lines show the Highest Posterior Density Intervals (HPDIs: 80%: thick; 95%: thin). Solid points indicate 95% HPDIs that exclude zero in the *Joint* (purple) and *Structured* (green) models.

**Supplemental Figure S2:**
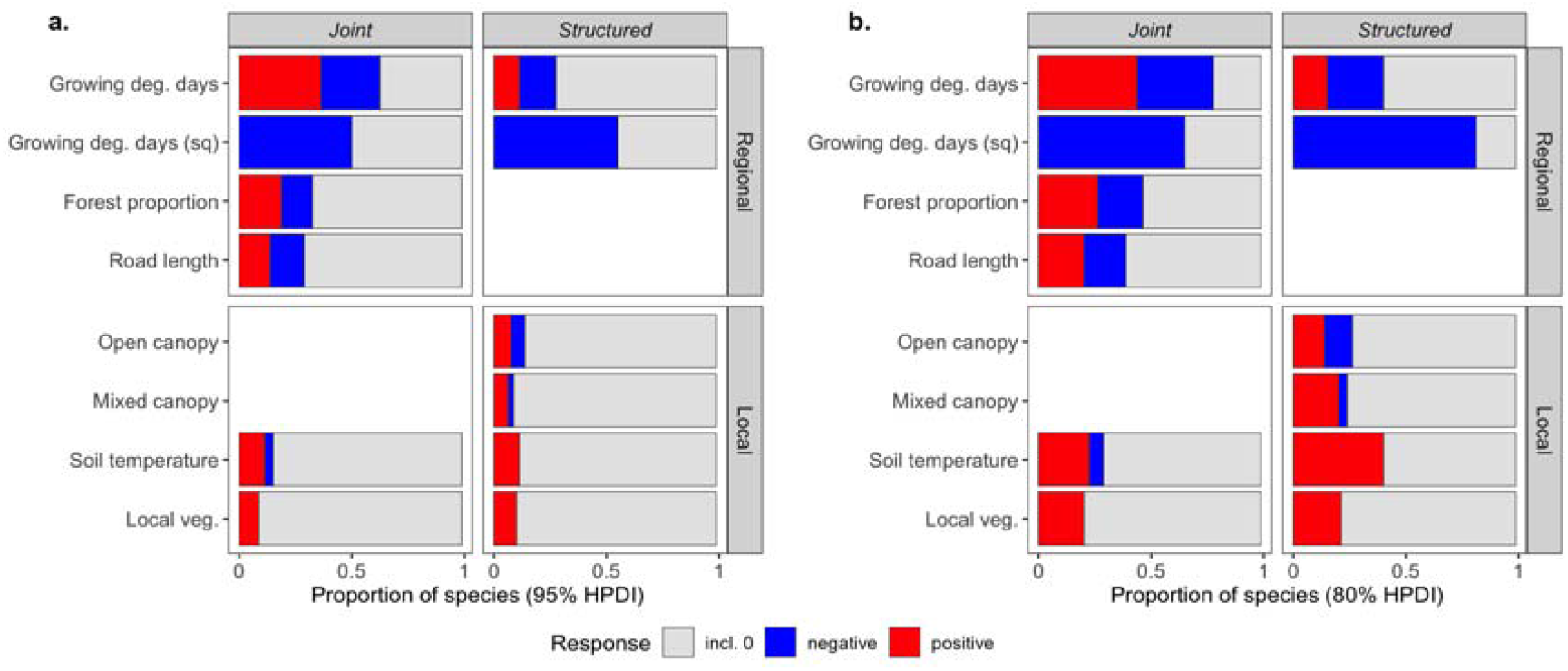
Proportion of species’ responses to each covariate. Responses are considered positive (red) or negative (blue) if the Highest Posterior Density Interval (HPDI) excludes zero for **(a)** 95% HPDIs and **(b)** 80% HPDIs.

**Supplemental Figure S2:**
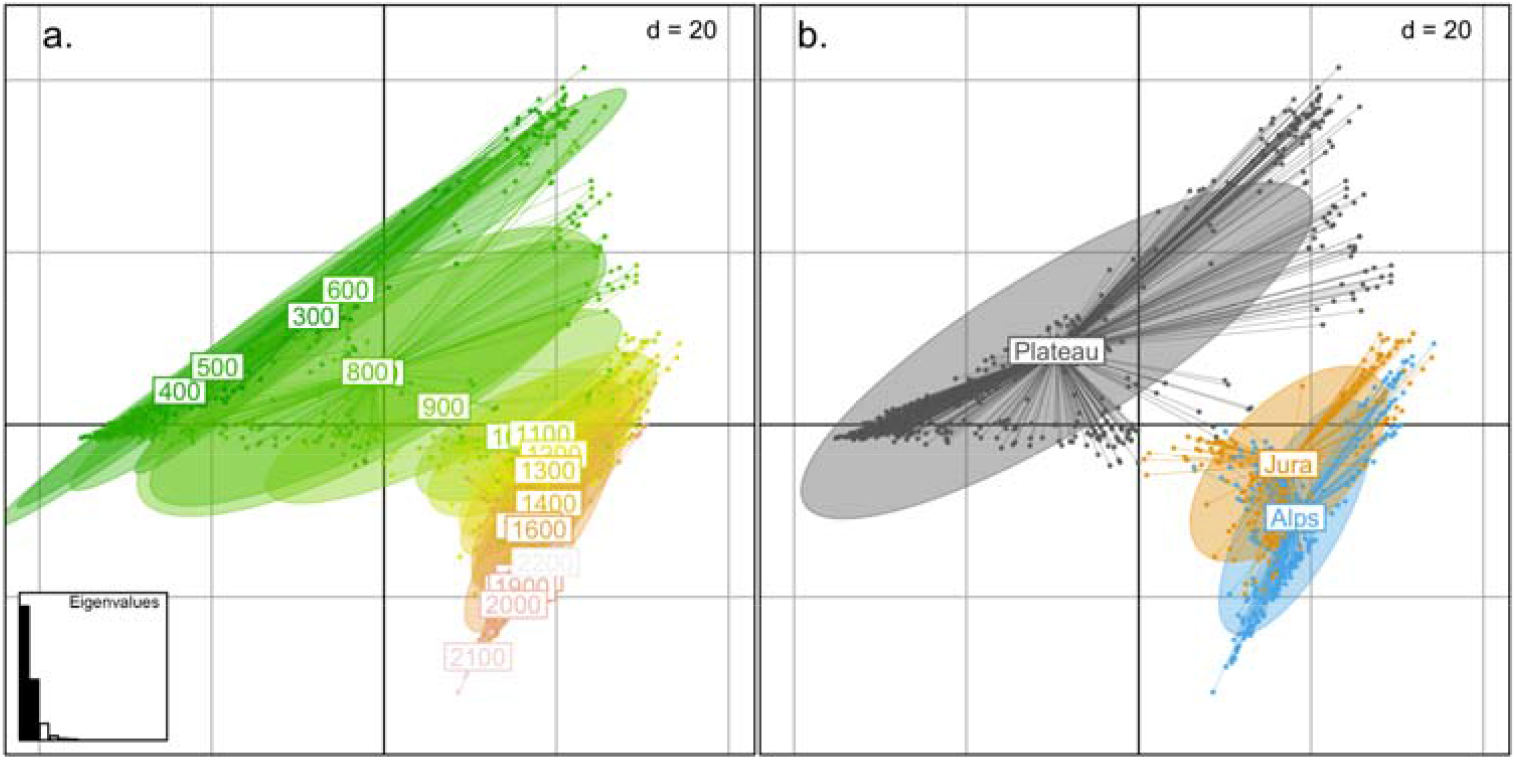
Double principle coordinate analysis (DPCoA) of local communities predicted by the *Structured* model, with plots colored by **(a)** elevational bin, and **(b)** region. The central plateau includes hills from 300m to 1000m, with the Jura and the Alps rising steeply in the east and west, respectively.

**Supplemental Figure S4:**
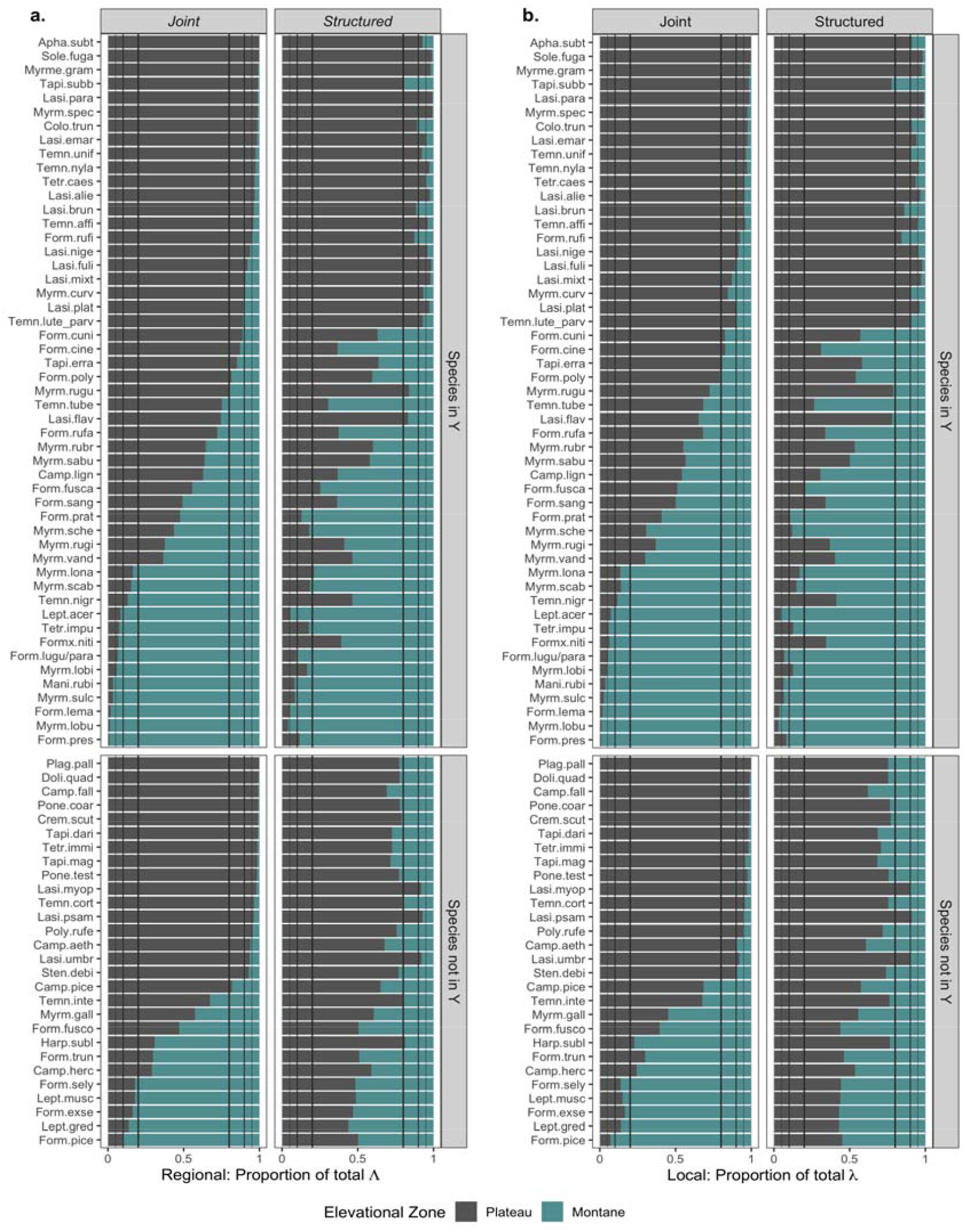
Posterior species intensity across plateau and montane zones in each model. Each species’ total intensity (a. Regional; b. Local) was partitioned into the proportion occurring in plateau (1000m: grey) and montane (1000m: green) elevations. Vertical lines indicate 80%, 90%, and 95% for either zone.

**Supplemental Figure S5:**
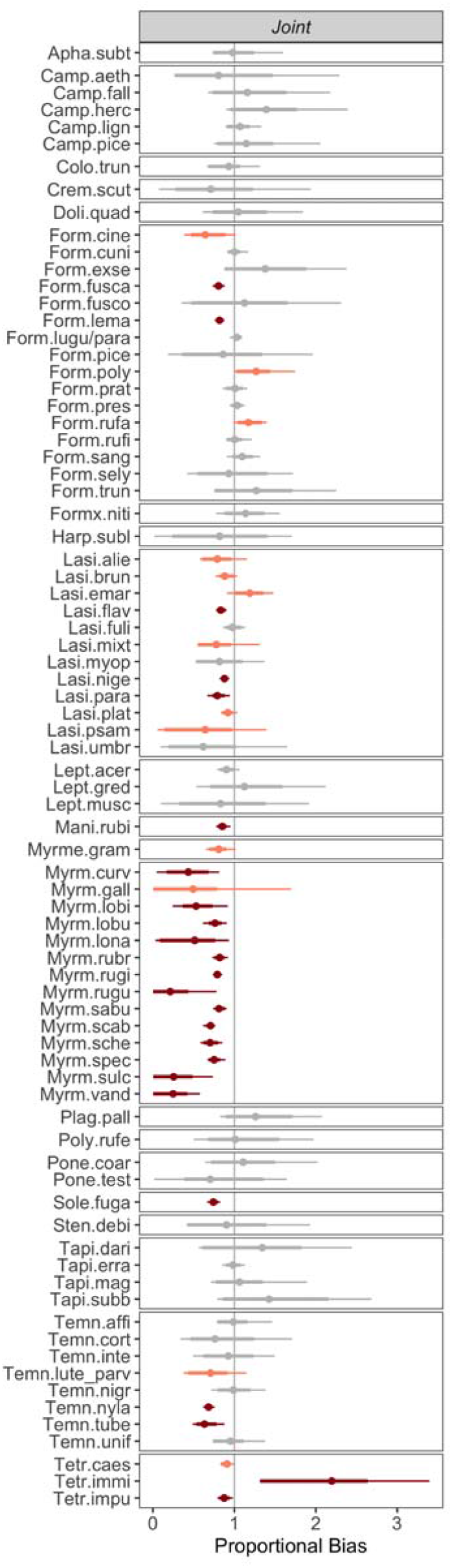
Proportional taxonomic bias posterior distributions. Medians and Highest Posterior Density Intervals (HPDIs; 80%: thick; 95%: thin) for the representation of each species in the presence-only dataset relative to the structured abundance dataset. Values less than 1 indicate under-representation in the presence-only dataset. Dark red: 95% HPDIs exclude zero; light red: 80% HPDIs exclude zero.

**Supplemental Figure S6:**
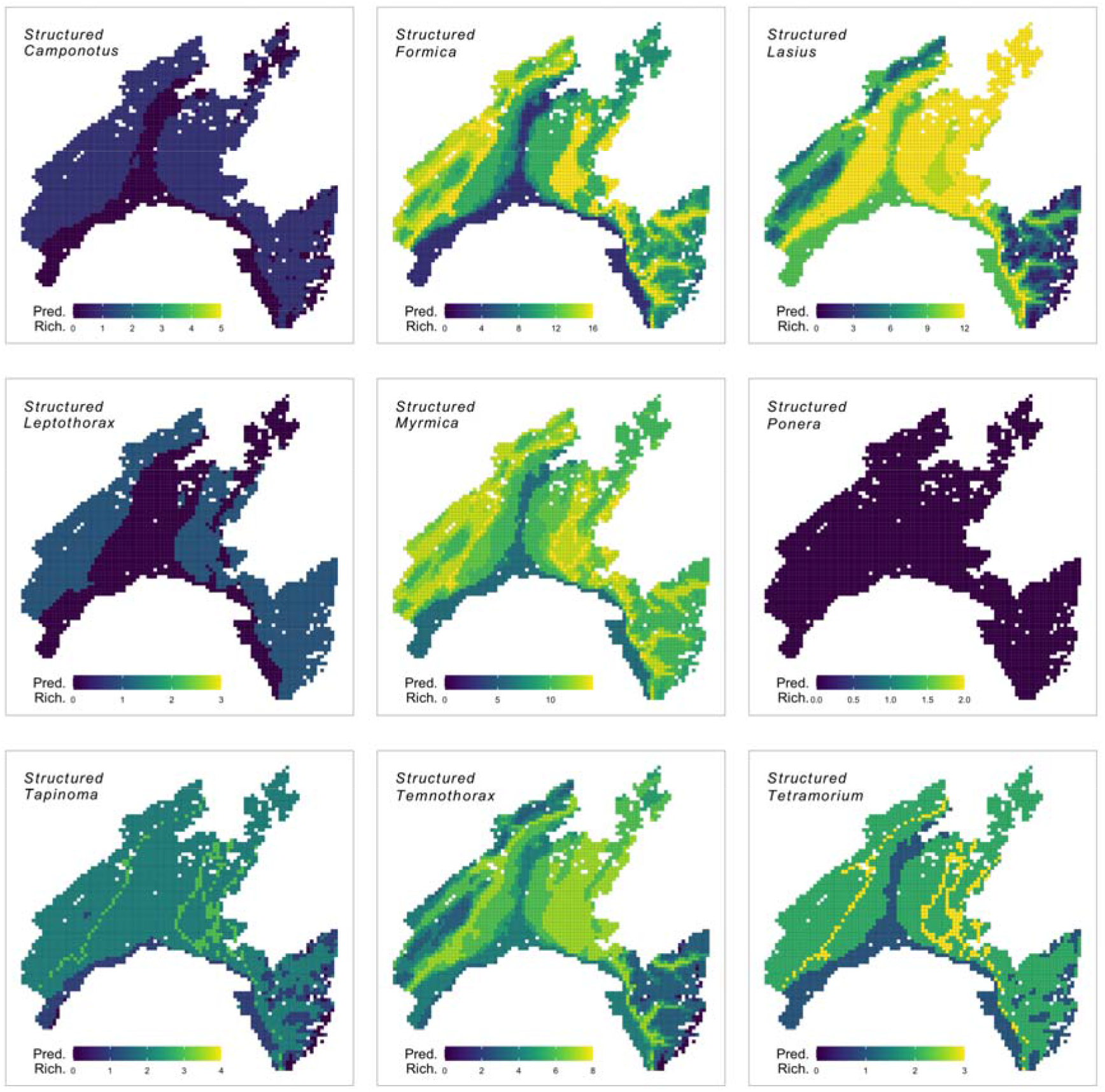
Maps of predicted richness in the *Structured* model for genera with at least two species. Predictions are based on 95% Highest Posterior Density Intervals for each species.

**Supplemental Figure S7:**
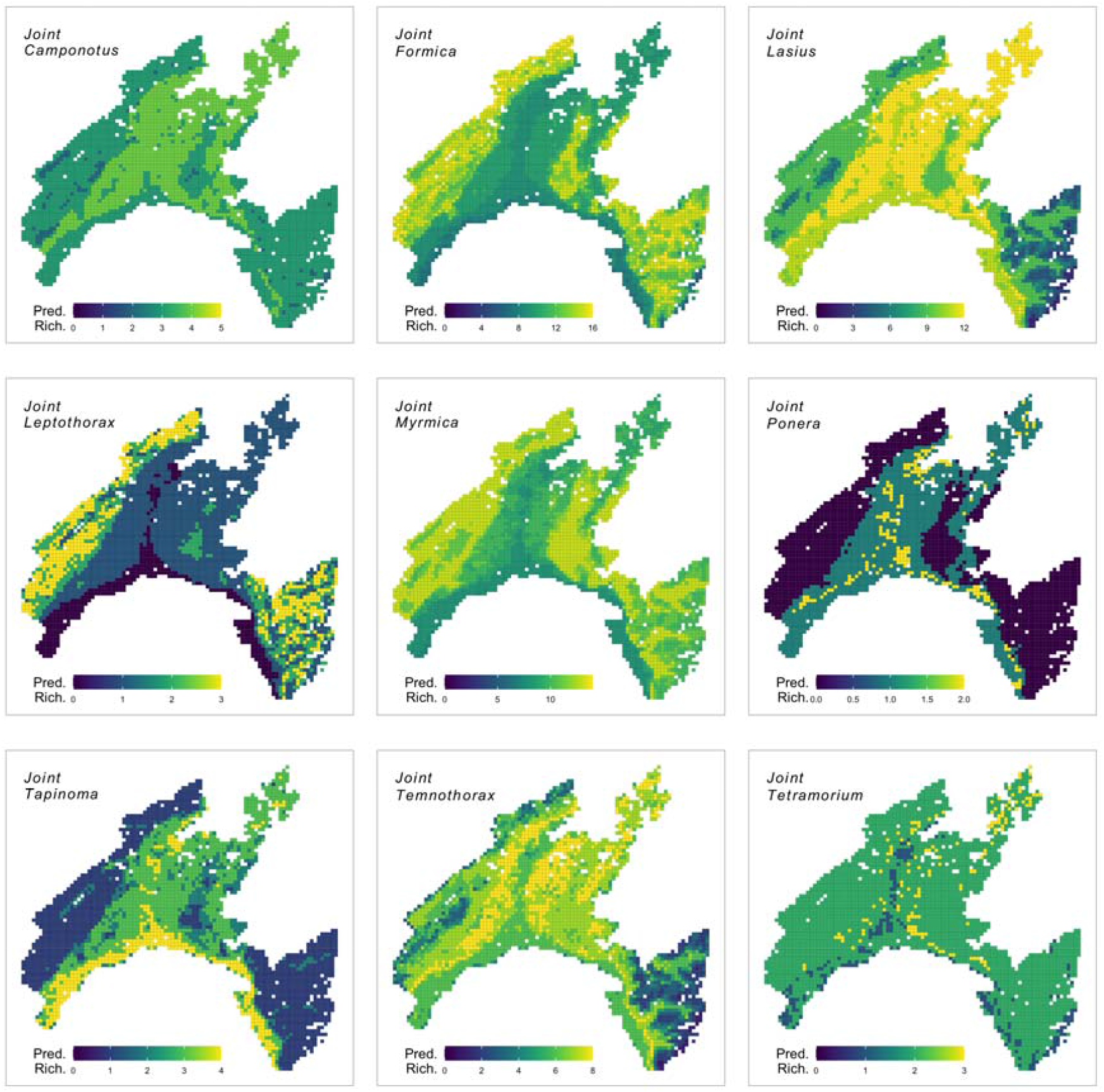
Maps of predicted richness in the *Joint* model for genera with at least two species. Predictions are based on 95% Highest Posterior Density Intervals for each species.

## Notes

https://github.com/Sz-Tim/opfo

https://github.com/Sz-Tim/5_citsci

https://github.com/glavanc1/ants_ID

### Competing Interest Statement

The authors have declared no competing interest.

